# Universal features of Nsp1-mediated translational shutdown by coronaviruses

**DOI:** 10.1101/2023.05.31.543022

**Authors:** Katharina Schubert, Evangelos D. Karousis, Ivo Ban, Christopher P. Lapointe, Marc Leibundgut, Emilie Bäumlin, Eric Kummerant, Alain Scaiola, Tanja Schönhut, Jana Ziegelmüller, Joseph D. Puglisi, Oliver Mühlemann, Nenad Ban

**Affiliations:** Department of Biology, Institute of Molecular Biology and Biophysics, ETH Zurich, Zurich, Switzerland; Department of Chemistry and Biochemistry, University of Bern, Bern, Switzerland; Department of Structural Biology, Stanford University School of Medicine, Stanford, CA, USA; Multidisciplinary Center for Infectious Diseases, University of Bern, Bern, Switzerland

## Abstract

Nonstructural protein 1 (Nsp1) produced by coronaviruses shuts down host protein synthesis in infected cells. The C-terminal domain of SARS-CoV-2 Nsp1 was shown to bind to the small ribosomal subunit to inhibit translation, but it is not clear whether this mechanism is broadly used by coronaviruses, whether the N-terminal domain of Nsp1 binds the ribosome, or how Nsp1 specifically permits translation of viral mRNAs. Here, we investigated Nsp1 from three representative *Betacoronaviruses* – SARS-CoV-2, MERS-CoV, and Bat-Hp-CoV – using structural, biophysical, and biochemical assays. We revealed a conserved mechanism of host translational shutdown across the three coronaviruses. We further demonstrated that the N-terminal domain of Bat-Hp-CoV Nsp1 binds to the decoding center of the 40S subunit, where it would prevent mRNA and eIF1A binding. Structure-based biochemical experiments identified a conserved role of these inhibitory interactions in all three coronaviruses and showed that the same regions of Nsp1 are responsible for the preferential translation of viral mRNAs. Our results provide a mechanistic framework to understand how *Betacoronaviruses* overcome translational inhibition to produce viral proteins.

## Introduction

Coronaviruses are enveloped, positive-stranded RNA viruses that infect animals and humans. Members of the *Betacoronavirus* (β-CoV) genus^1^ cause serious respiratory diseases in humans: severe acute respiratory syndrome coronavirus 2 (SARS-CoV-2) caused the COVID-19 pandemic^2, 3^, and SARS-CoV and Middle East respiratory syndrome coronavirus (MERS-CoV) were responsible for two previous epidemics^4^. The approximately 30 kb β-CoV genome starts with a capped 5’ untranslated region (5’UTR), contains many protein-encoding open reading frames, and ends with a polyadenylated 3’UTR^5^. The first sixteen viral proteins produced are nonstructural proteins (Nsps). Many Nsps work collectively to render cellular conditions favorable for viral infection through unclear mechanisms, whose definition could accelerate the development of new therapeutics.

Nsp1, the first viral protein synthesized, is a major pathogenicity factor of SARS-CoV-2. It inhibits host protein synthesis, induces host mRNA cleavage^6^, and plays a role in mRNA export and immune evasion pathways^7,8,9,10,11,12^. How Nsp1 inhibits host translation is well established and supported by structures of the human small ribosomal subunit (40S) bound by Nsp1 from SARS-CoV-2^9, 13, 14^. The C-terminal domain (CTD) of Nsp1 binds to the entry region of the mRNA channel on the 40S subunit, where it sterically clashes with mRNA^15^, thereby inhibiting translation. Whether this mechanism extends to Nsp1 proteins from other β-CoVs remains unclear.

The role of the Nsp1 N-terminal Domain (NTD) is enigmatic. Biochemical data indicate that SARS-CoV-2 genomic and sub-genomic RNAs overcome Nsp1-mediated inhibition to produce viral proteins^6, 16–19^. The Nsp1 NTD was suggested to play a crucial role in this process: mutation of R124A/K125A and R99A in this domain prevents preferential translation of viral mRNAs^6^. Viral protein production also relies on distinct nucleotides (C15, C19, and C20) within the first stem-loop (SL1) of the viral mRNAs, referred to as the leader sequence^20^. However, the structural basis of how the Nsp1 NTD selectively allows viral mRNAs to form productive complexes with the 40S subunit during translation initiation is unclear, as this domain has never been visualized in ribosome-Nsp1 complexes^21,22,23^.

We sought to better understand how β-CoV Nsp1 proteins inhibit host translation and how coronaviruses overcome such inhibition to produce viral proteins. We selected Nsp1 proteins from three representative β-CoVs, which included SARS-CoV-2 (subgenus: *Sarbecovirus*). We also studied Nsp1 proteins from MERS-CoV (subgenus: *Merbecovirus*) and a β-CoV that infects bats, Bat-Hp-CoV_Zhejiang2013 (referred to as Bat-Hp-CoV hereafter). Bat-Hp-CoV was included as it is the only member of the *Hibecovirus* subgenus, and its Nsp1 protein was shown to bind the human ribosome^15^. Furthermore, all known human β-CoVs likely originate from viruses that originally infect animals^4^, with bats and rodents being the dominant source for transmission via intermediate hosts, such as camels for MERS-CoV and possibly civets/pangolins/bats for SARS-CoV and SARS-CoV-2^24, 25^ (Figure 1). In this study, we provide structural and biochemical evidence that Nsp1 from all three tested β-CoVs suppresses the translation of host mRNAs by binding to the mRNA channel of the 40S ribosomal subunit. Using paired structural and single-molecule analyses, we further reveal that the NTD of Bat-Hp-CoV Nsp1 binds to the decoding center of the 40S subunit, a functionally critical region of the ribosome. Our results rationalize several previous and new biochemical observations^6, 16, 18^, and they suggest a conserved mechanism for how Nsp1 in β-CoVs selectively inhibits translation of cellular mRNAs.

**Figure 1:**
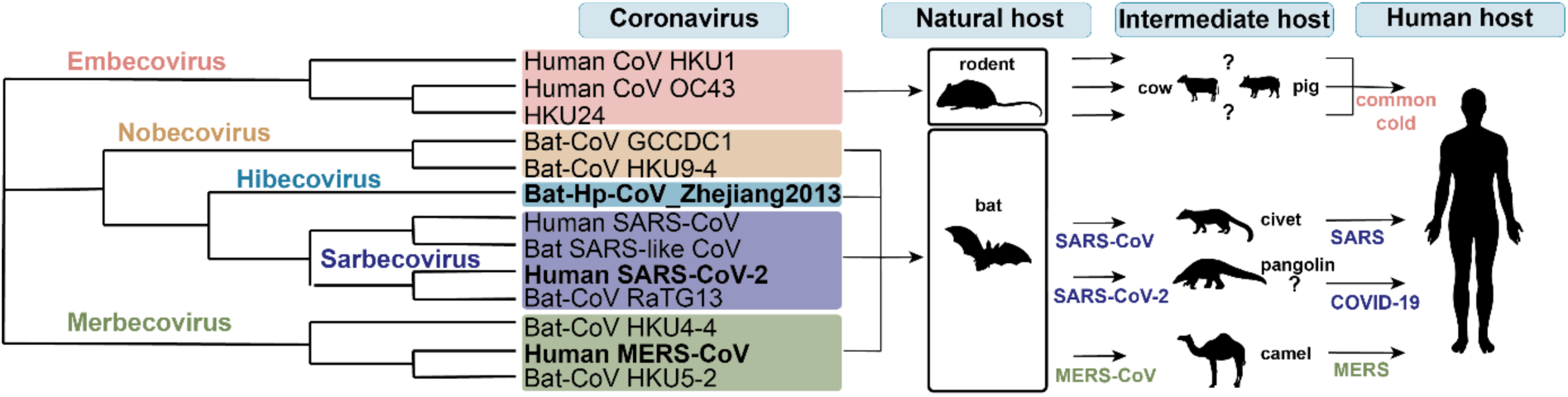
Phylogenetic tree representing the evolutionary relationships among selected β-CoVs. Out of the five β-CoV subgenera, *Embeco-, Nobeco-, Hibeco-, Sarbeco*- and *Merbecovirus*, rodents are the natural reservoir for members of *Embecoviruses* and bats for the other four subgenera. Via intermediate hosts, including civets, pigs, and camels, several CoVs have adapted to also infect humans, causing common cold symptoms and more serious diseases such as SARS, MERS, and COVID-19. The viruses investigated in this manuscript are highlighted in bold.

## Results

### MERS-CoV Nsp1 binds the 40S ribosomal subunit to inhibit translation

Similar to other β-CoVs, MERS-CoV Nsp1 inhibits host gene expression^26^. However, due to sequence differences in its CTD relative to SARS-CoV-2 Nsp1, it has been proposed that MERS-CoV Nsp1 acts independently of the human ribosome and selectively targets host mRNAs in the nucleus^26^.

To identify and functionally characterize cellular complexes that interact with MERS-CoV Nsp1, we combined biochemical and structural analyses. FLAG-tagged MERS-CoV Nsp1 was transiently expressed in HEK293E cells, affinity purified and subjected to ultracentrifugation to isolate large molecular weight complexes. Purified complexes were analyzed using high-resolution cryo-EM. We identified the 43S pre-initiation complex (PIC) as the main component of the sample and determined its structure at 2.65 Å resolution (Figure S1). Despite its divergent sequence (Figure S3B), we found the CTD of MERS-CoV Nsp1 bound to the mRNA entry channel of the 40S subunit as observed in the SARS-CoV-2 complex (Figure 2A). Eukaryotic initiation factors (eIF) eIF1 and eIF1A reside at the 40S ribosomal P- and A-sites, respectively^9, 13^. The complex also contained eIF2αβγ-tRNA_i_ and all subunits of eIF3^9^ except the eIF3j subunit (Figure S2), consistent with prior biochemical and structural experiments^9, 13, 15^. Our well-resolved cryo-EM maps permitted visualization and interpretation of several previously unseen or unassigned regions of the eIFs, allowing the generation of a more complete structural description of the human 43S PIC complex (Figure S2).

**Figure 2:**
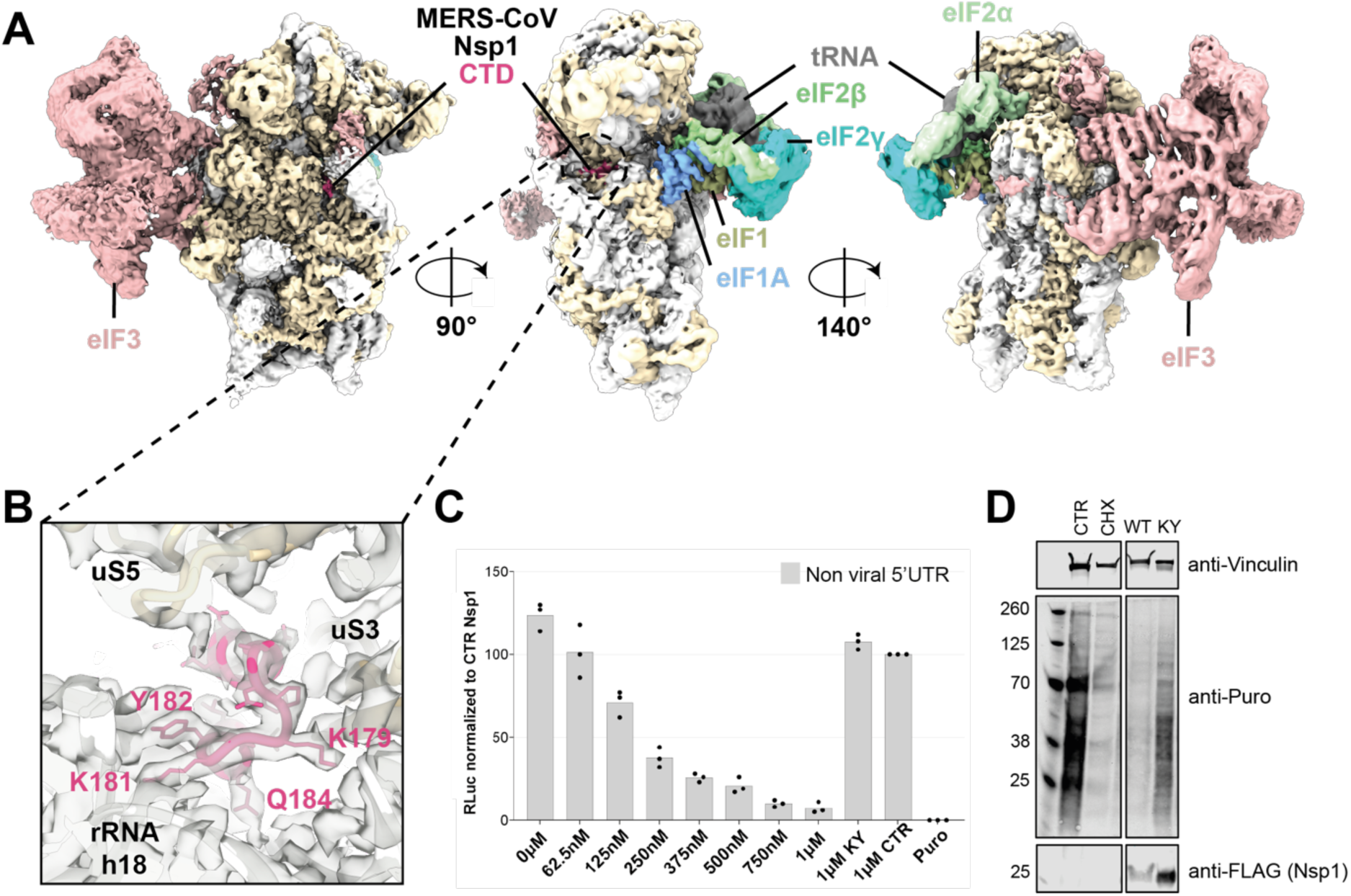
The C-terminal domain of MERS-CoV Nsp1 binds to the 43S PIC as observed for SARS-CoV-2 Nsp1 and inhibits translation. **(A)** Views of MERS-CoV Nsp1 (CTD in magenta) binding to the small ribosomal subunit (rRNA in light grey, ribosomal proteins in beige) with initiation factors eIF1 (light green), eIF1A (light blue), eIF3 (rose brown), eIF2αβγ (different greens), and P-site initiator tRNA (grey). The map is filtered to 5 Å. **(B)** Close-up view of MERS-CoV Nsp1 CTD binding to the mRNA entry channel (map filtered to local resolution). **(C)** Relative *Renilla* luciferase (RLuc) activity measurements of *in vitro* translation reactions with increasing amounts of purified WT MERS-CoV Nsp1 or the KY mutant, normalized to the reaction in the presence of the control protein (CTR) (Trx1-SSG-CTD KH SARS-CoV-2, see methods). Data are presented as mean values of three biological replicates (sets of translation reactions) averaged after three measurements. The negative control reaction contained 320 μg/ml puromycin (Puro). **(D)** Western blot of a puromycin incorporation assay in the presence of Nsp1. HEK293 cells were transfected with MERS-CoV wild-type FLAG-tagged Nsp1 (WT) or FLAG-tagged Nsp1 with a mutated KY motif (KY). Cells transfected with an empty vector served as a control in the DMSO and CHX reactions. As a control, two cultures were transfected with an empty plasmid and incubated with cycloheximide (CHX) or DMSO (CTR). Before collection, nascent proteins were labeled during a puromycin pulse. Cell lysates were prepared, and the Nsp1 expression (anti-FLAG) and incorporated puromycin (anti-Puro) were analyzed by Western blotting. Vinculin served as a loading control.

The CTD of MERS-CoV Nsp1 binds to the mRNA channel of the 40S subunit, where it would interfere with mRNA binding and forms a network of molecular interactions analogous to SARS-CoV-2 Nsp1 (Figure 2B). In place of the highly conserved _164_KH_165_ motif in SARS-CoV-2, the corresponding K181 and Y182 residues in MERS-CoV Nsp1 mediate ionic and stacking interactions with U607 and U630 of the 18S rRNA, respectively (Figure S3A). The second α-helix in the CTD is shorter than in SARS-CoV-2 Nsp1; however, the contacts mediated by the essential positively charged amino acid residues in SARS-CoV-2, R171, and R175, are replaced by two lysine residues (K188 and K189) in MERS-CoV (Figure S3A, B).

We verified our structural findings using a human-based *in vitro* translation assay, which revealed a concentration-dependent inhibition of translation upon adding wild-type (WT) MERS-CoV Nsp1 protein (Figure 2C). The substitution of residues K181 and Y182 with alanine (KY mutant) alleviated Nsp1-dependent inhibition of translation (Figure 2C). Therefore, the binding mode of Nsp1 to the 40S subunit and the resulting translation inhibition activity is preserved between SARS-CoV-2 and MERS-CoV, including the KY motif in MERS-CoV that functionally replaces the KH motif of SARS-CoV-2 and is equally critical for 40S binding.

Since our *in vitro* data indicated that MERS-CoV Nsp1 inhibits translation, we examined translation inhibition *in vivo*. To determine whether MERS-CoV Nsp1 inhibits human translation in HEK293 cells, we transiently expressed FLAG-tagged WT Nsp1 or the KY CTD mutant, which cannot bind to the mRNA channel, and assessed the effect on global translation using a puromycin incorporation assay. Compared to cells transfected with a control plasmid, MERS-CoV WT Nsp1 reduced puromycin incorporation and translation efficiency (Figure 2D), whereas the KY mutant did not. These results corroborate our *in vitro* translation assays and highlight the importance of the KH and KY motifs in the CTD of Nsp1 across β-CoVs for ribosome binding and translation inhibition^9, 13, 14^.

### Nsp1 inhibits translation by binding to both the mRNA channel and the A-site of the 40S

To further investigate the mechanism of host translation shutdown across representative examples of *Betacoronavirus* species, we also determined the high-resolution cryo-EM structure of the Bat-Hp-CoV Nsp1 in complex with the 40S subunit using an *in vitro* reconstitution approach. In addition to the Bat-Hp-CoV Nsp1 CTD bound in the mRNA entry channel, we resolved for the first time the density for the Nsp1 linker and NTD regions in about 15% of 40S complexes (Figure S4). Based on these results, and considering previous binding affinity measurements indicating cooperative binding of SARS-CoV-2 Nsp1 with eIF1^15^, we reconstituted the complex with both Bat-Hp-CoV Nsp1 and eIF1 bound to the 40S subunit to obtain a more complete overview of how Nsp1 interacts with the 40S subunit during initiation, (Figure 3, Figure S5).

**Figure 3:**
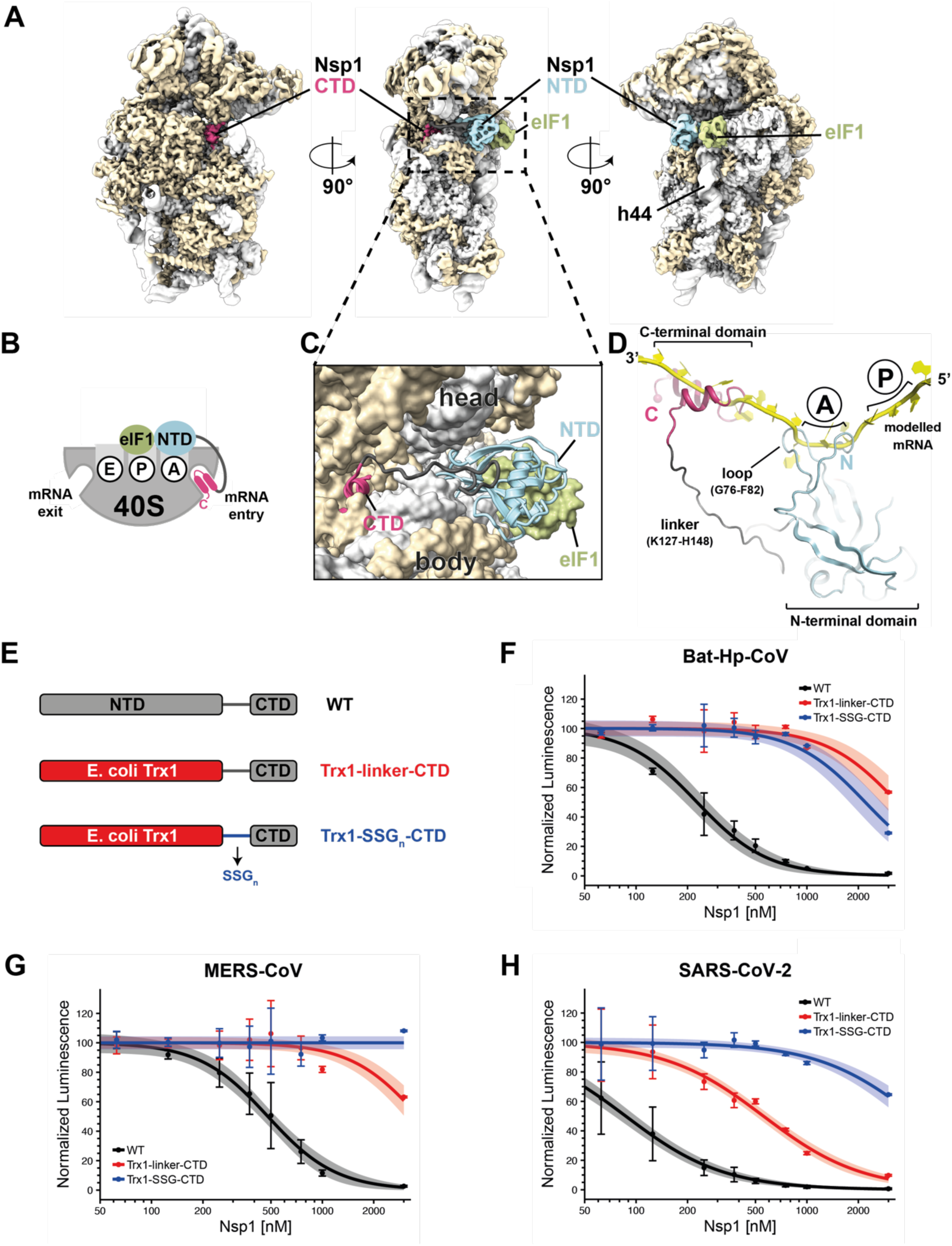
Binding interfaces of both the CTD and NTD of Bat-Hp-CoV Nsp1 visualized on the small ribosomal subunit by determining the cryo-EM structure of an *in vitro* reconstituted complex. **(A)** Views of Bat-Hp-CoV Nsp1 (CTD in magenta, linker in dark grey and NTD in light blue) binding to the small ribosomal subunit (rRNA in light grey, ribosomal proteins in beige) with initiation factor eIF1 (light green) bound to the ribosomal P-site. The map is filtered to local resolution. **(B)** Schematic overview of the Nsp1 domain arrangement in the ribosomal complex, including the Nsp1 NTD (light blue), a flexible linker (dark grey), Nsp1 CTD (magenta), and eIF1 (light green). **(C)** Close-up model view of Bat-Hp-CoV Nsp1 binding to the mRNA entry channel and decoding site (ribosomal A-site). **(D)** Superposition of a canonically bound mRNA (yellow) (PDB 7O7Z) reveals that Nsp1 NTD contributes to preventing classical binding of mRNA at the A-site. **(E)** Schematic of Nsp1 WT and with the NTD mutated to *E. coli* Trx1 and the linker mutated to repetitive SSG modules. **(F-H)** Relative RLuc activity measurements of *in vitro* translation reactions with increasing amounts of purified Bat-Hp-CoV, SARS-CoV-2 and MERS-CoV Nsp1 WT, Trx1-linker-CTD or Trx1-SSG-CTD. Data are presented as mean values ± standard deviations of three biological replicates (sets of translation reactions) averaged after three measurements and were normalized to 0 nM Nsp1. The data were fit using a dose-response model, requiring the same Hill coefficient for WT, Trx1-linker-CTD, and Trx1-SSG-CTD and the shaded area corresponds to the 95% confidence interval. Data for MERS-CoV Trx-SSG-CTD was fit using an intercept model. Data was rescaled such that the intercept of all fits was equal to 100 normalized luminescence.

The 3.0 Å structure of the Bat-Hp-CoV Nsp1 – 40S – eIF1 complex revealed density for the entire Nsp1 protein and allowed us to visualize the NTD bound to the decoding center of the 40S subunit. Originating from the CTD bound within the mRNA channel (Figure 3A, Figure S3A), the approximately 20-amino acid long linker region, considered flexible in solution^21, 23^, interacts with uS3 in the head of the 40S subunit and extends towards the decoding center, where the NTD binds. The NTD adopts the same fold as seen previously for the isolated NTD from SARS-CoV-2^22, 23^ and occupies the site on the 40S subunit where eIF1A resides in canonical initiation complexes (Figure 3B, C) (for details, see next chapter). Despite evidence for cooperative binding, we do not observe direct interactions between Nsp1 NTD and eIF1, as is the case for eIF1A and eIF1 in initiation complexes^27, 28^. The cooperative binding is likely mediated through independent stabilization of the rRNA helix 44 (h44) conformation, to which both Nsp1 and eIF1 bind^27, 28^. Two loops in the NTD of Nsp1 enter the mRNA channel on the 40S subunit and would clash with a canonically bound mRNA. Therefore, both the CTD and the NTD of Bat-Hp-CoV Nsp1 prevent mRNA binding on the entrance side of the mRNA channel and in the decoding center of the 40S subunit, respectively (Figure 3D).

To assess the inhibitory effect of the NTD and the linker on translation, as suggested by the structure, we replaced the Nsp1 NTD with bacterial thioredoxin (*E. coli* Trx1) only, and in another construct, in addition to the NTD, also the linker with a repetitive SSG sequence and performed *in vitro* translation assays (Figure 3E). For all three tested β-CoVs, the substitution of the NTD reduced the inhibitory activity of Nsp1. Compared to WT Nsp1, the concentration of Trx1-carrying Nsp1 mutants with the WT linker needed to reduce translation to 50% (apparent dissociation constant K_D_) increased by more than an order of magnitude for Bat-Hp-CoV and MERS-CoV, whereas a ∼6-fold higher concentration of SARS-CoV-2 was required (Figures 3F-H). The latter aligns with the high protein concentrations (6 μM) required in earlier analogous assays for SARS-CoV-2 Nsp1^13^. In agreement with our structural findings, we observed that also the linker contributes to the Nsp1 translation inhibition capacity and shifts the apparent K_D_ even further to higher concentrations, which is particularly pronounced for SARS-CoV-2 (Figure 3H). Our findings indicate that the CTD interaction with the mRNA entry channel of the 40S subunit largely provides the affinity for Nsp1 binding, as mutations in the CTD abolished translation inhibition activity (Figure 2C)^9, 13^. However, our titration experiments demonstrate that the NTD plays an important role in translation inhibition at lower Nsp1 concentrations, like those that are most likely reached during the ramp-up of viral translation in infected cells.

### The Nsp1 interaction with the 40S decoding center is specific and dynamic

The full-length Bat-Hp-CoV Nsp1 structure in complex with the 40S subunit revealed the specificity of interactions between the NTD and the decoding center. The Nsp1 NTD of Bat-Hp-CoV interacts at multiple sites with the 40S subunit, including h18 and the two decoding bases in rRNA helix h44 (Figure 4). A conserved region in the NTD, _97_GRSG_100_ (corresponding to _98_GRSG_101_ in SARS-CoV-2), stabilizes both decoding nucleotides (A1824 and A1825) in h44 in a flipped-out conformation (Figure 4C, D and Figure S6). Interestingly, in canonical 43S PIC and MERS-CoV Nsp1 43S PIC, eIF1A stabilizes only A1825 in a flipped-out conformation, while in non-initiating, factor-less 40S subunits, both decoding bases are disordered (Figure S6). Previous studies showed that mutating R99 in SARS-CoV-2 within this motif prevented the escape of SARS-CoV-2 viral mRNAs from translational shutdown^6^. Interestingly, despite the low sequence conservation of MERS-CoV compared to SARS-CoV-2 (approximately 21% sequence identity^29^) and Bat-Hp-CoV (approximately 26%), the second glycine in the motif is conserved among these viruses. When modeling the MERS-CoV NTD or superimposing the structure of the SARS-CoV-2 NTD^22, 23^, the corresponding glycines would be positioned at the same location as residue G100 in Bat-Hp-CoV Nsp1 in the 40S complex, which argues for a conserved role in interactions with the decoding bases. The second prominent interaction site observed between Bat-Hp-CoV Nsp1 and the 40S subunit includes another conserved region in the NTD (_122_LRRRG_126_) just before the linker region. The conserved arginine R123 (corresponding to R124 in SARS-CoV-2 and R146 in MERS-CoV) interacts with Um627 in h18 (Figure 4C, E). Mutations of this arginine in SARS-CoV-2 also prevented selective translation of viral mRNAs^6^.

**Figure 4:**
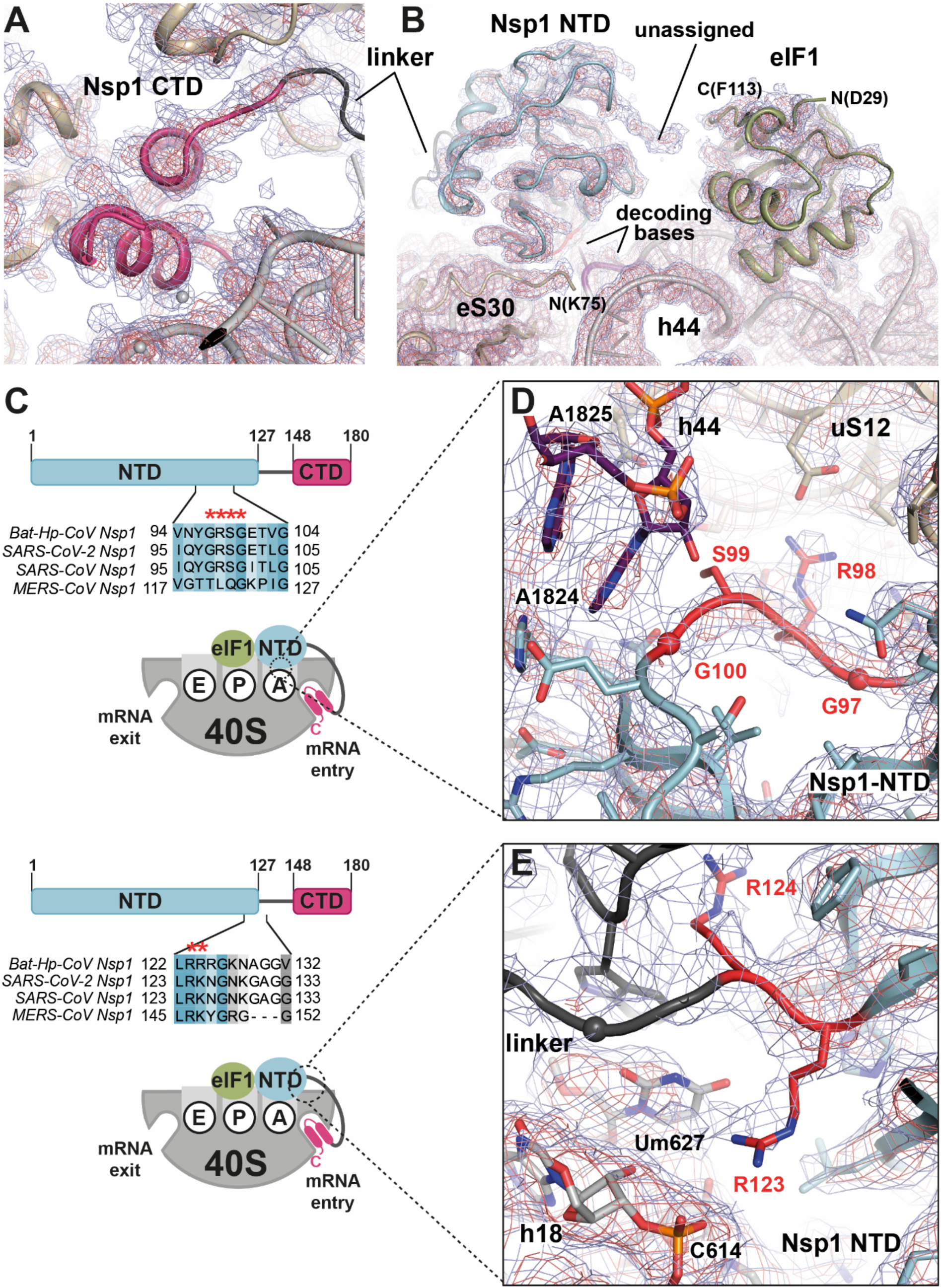
Details of Bat-Hp-CoV Nsp1 binding to the small ribosomal subunit. **(A-B)** Overview of Bat-Hp-CoV Nsp1 CTD (magenta) binding to uS3, uS5 (beige), and the 18S rRNA (grey) in the mRNA entry channel as well as Nsp1 NTD (light blue) and eIF1 (light green) binding at the intersubunit space. The 3.0 Å experimental EM densities are contoured at 7σ (blue) and 11σ (red), respectively. The unassigned density likely originates from one of the ribosomal proteins located in the head of the 40S. **(C)** Clustal Omega sequence alignment of Nsp1 from Bat-Hp-CoV, SARS-CoV-2, SARS-CoV, and MERS-CoV, colored according to conservation using Jalview^30^, with conserved residues mutated in this study, indicated by red asterisks. The schematics indicate the zoomed positions shown in (D) and (E) and are color-coded as in (C). **(D)** Detailed view of the decoding site with Nsp1 NTD (light blue) bound to the A-site. In the presence of Nsp1, both decoding nucleotides A1824 and A1825 (purple) in h44 adopt a flipped-out conformation. The conserved _97_GRSG_100_ motif of Nsp1 (numbering according to Bat-Hp-CoV) is depicted in red. **(E)** Detailed view of Nsp1 interaction with h18. Nsp1 NTD is depicted in light blue, the conserved region _123_RR_124_ in red, and the linker region in dark grey. The 3.0 Å experimental EM densities shown as blue and red meshes are contoured at 6σ and 9σ, respectively.

Enabled by our structure, we devised single-molecule experiments to examine the dynamics of Nsp1 association with the decoding center. The proximity of the N-terminus of Bat-Hp-CoV Nsp1 to an established^15^ 40S subunit labelling site (N-terminus of uS19) (∼85 Å) might provide a Förster Resonance Energy Transfer (FRET) signal that reports on decoding center occupancy (Figure 5A). To test this hypothesis, we labelled the N-terminus of both SARS-CoV-2 and Bat-Hp-CoV Nsp1 proteins with Cyanine-5 dye (Cy5) (FRET acceptor). A molar excess of Cy5-labeled Nsp1 proteins was incubated with Cy3-labeled 40S subunits (FRET donor) and excess unlabeled eIF1. Nsp1 – 40S – eIF1 complexes were then bound to a 5’-biotinylated RNA that associates with the 40S subunit outside the decoding center and mRNA entry channel. Final complexes were tethered specifically to neutravidin-coated surfaces and imaged at equilibrium using single-molecule spectroscopy (Figure S7A, B).

**Figure 5:**
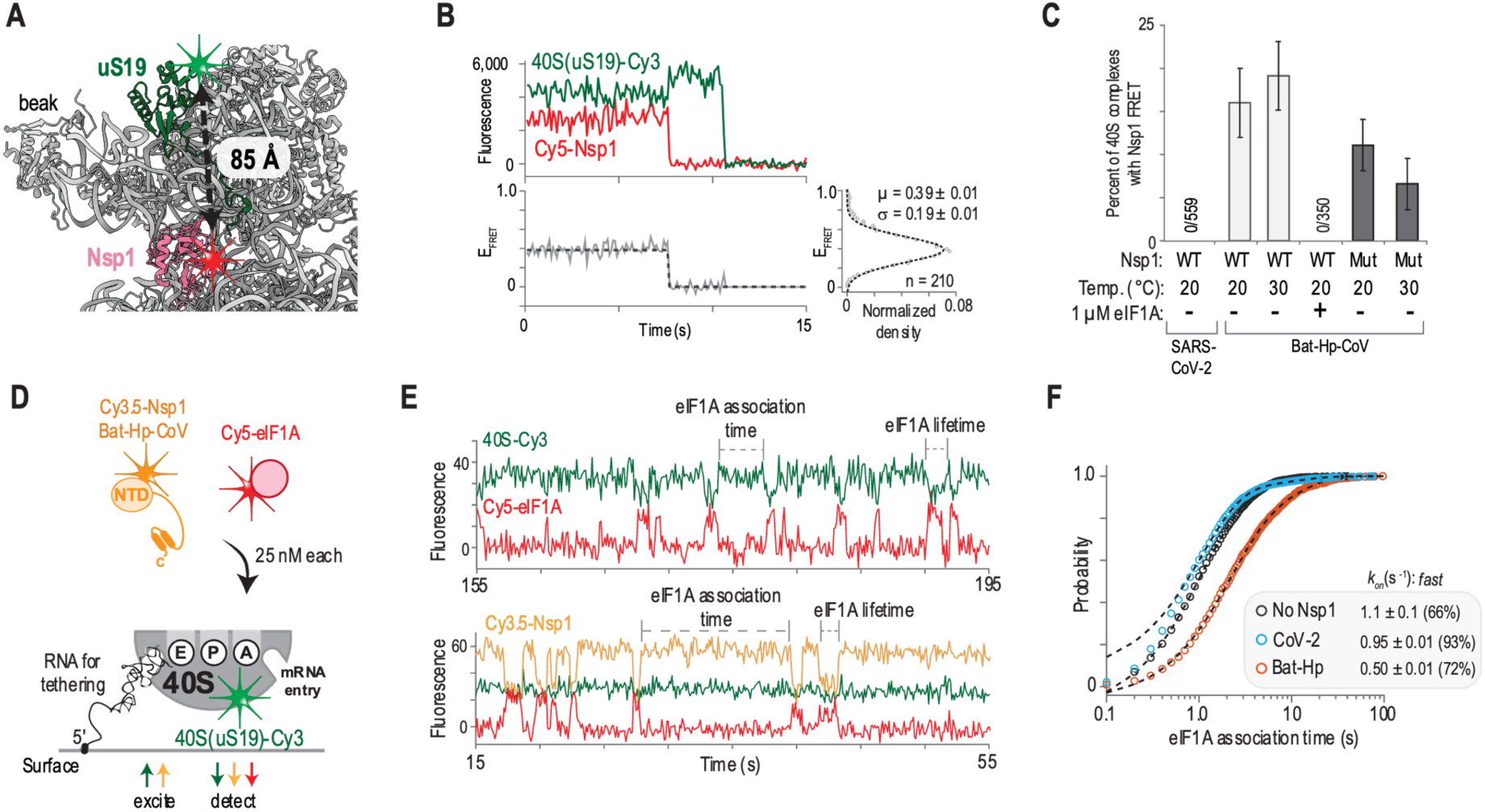
The NTD of Bat-Hp-CoV Nsp1 stably docks within the decoding center of the 40S subunit. **(A)** The FRET labels in the uS19 N-terminus and the Bat-Hp-CoV NTD Nsp1 are ∼85 Å apart when the Nsp1 NTD is bound to the decoding center of the 40S subunit, as observed in our structure. The 40S subunit was labeled with Cy3 dye (FRET donor, green), and Nsp1 with Cy5 dye (FRET acceptor, red). **(B)** Example of single-molecule fluorescence data from a single ZMW for the Bat-Hp-CoV Nsp1(Cy5)-40S(Cy3) complex imaged at equilibrium at 20°C using total internal reflection fluorescence microscopy (TIRFm). **(C)** Quantification of the fraction of tethered 40S complexes (scored via the 40S(Cy3) signal, in either TIRFm or ZMW-based imaging) that yielded FRET. Under the indicated reaction conditions, measurements were either performed with wild-type (WT) Nsp1 or a mutant (Mut) carrying a GRSG-AAAA substitution. **(D)** Schematic of the real-time competition experiment (ZMW-based) that examined Nsp1 and eIF1A association with the 40S subunit simultaneously. The 40S subunit was labeled with Cy3 (green), Nsp1 with Cy3.5 (orange), and eIF1A with Cy5 (red). The presence of the 40S subunit and Nsp1 was detected via direct excitation using a 532 nm laser, with eIF1A occupancy revealed by FRET with either the 40S subunit or Nsp1. **(E)** Example of single-molecule fluorescence data with eIF1A lifetime (duration of eIF1A-Cy5 events) and association times (dwell between eIF1A-Cy5 binding events) indicated. **(F)** Quantification of eIF1A (25 nM final concentration) association times when the Nsp1 proteins (25 nM) were present, as indicated. Dashed lines represent fits to exponential functions.

Consistent with our structural results, we observed low-efficiency FRET (mean ∼0.4) between Bat-Hp-CoV Nsp1 and the 40S subunit on a subset of complexes (10-20%) (Figure 5B, C and Figure S7C). The observed FRET signal persisted for at least 100 s (k_off_ < 0.01 s^-1^) (Figure S7D-F) and was eliminated in the presence of excess eIF1A (Figure 5C), which binds competitively to the decoding center of the 40S subunit. Thus, the FRET signal indicated Nsp1 occupancy at the 40S decoding center. This interaction was destabilized by alanine substitution of the Nsp1 GRSG motif, which directly contacts the decoding bases; the number of complexes that yielded FRET decreased and they had modestly decreased lifetimes, particularly at an increased temperature (Figure 5C and Figure S7F). In contrast to the Bat-Hp-CoV protein, SARS-CoV-2 Nsp1 failed to yield FRET with the 40S subunit (0/559 complexes) (Figure 5C and Figure S7C). To explore the decoding-center interaction in more detail, we also performed competition experiments where association of Bat-Hp-CoV Nsp1 and eIF1A with the tethered 40S subunit was monitored in real-time (Figure 5D, E). Consistent with our findings above, the presence of Bat-Hp-CoV Nsp1 on the 40S subunit inhibited eIF1A association by at least 2-fold (Figure 5F), without impacting the lifetime of eIF1A on the complex (Figure S7G, H). eIF1A binding kinetics were unaltered by SARS-CoV-2 Nsp1 (Figure 5F and Figure S7G, H). This result further demonstrates that the NTD of SARS-CoV-2 Nsp1 cannot dock with a long residence time in the 40S decoding center. Thus, our single-molecule and structural findings indicate a conserved, bi-partite interaction occurs between Nsp1 and the 40S subunit, with variable binding kinetics across the three examined β-CoVs (see Discussion).

### Decoding center interacting regions of Nsp1 are important for the selective translation of viral mRNAs

To investigate the role of specific residues within the Nsp1 NTD that contact the 40S decoding center, we first tested in translation lysates and human cells whether the function of Nsp1 in the selective translation of viral mRNAs is conserved across the three β-CoVs. We generated reporter mRNAs that contained the 5’UTRs from SARS-CoV-2, MERS-CoV, or Bat-Hp-CoV, followed by the *Renilla* luciferase open-reading frame (Figure 6A). We assessed the *in vitro* translation output in the presence or absence of Nsp1 proteins at a concentration of 300 nM, a condition where the N-terminal domain plays an important role in translation inhibition as indicated by our biochemical experiments described above (Figure 3F-H). Notably, our assay recapitulated previous reports^17^ that the SARS-CoV-2 reporter mRNA evaded Nsp1-mediated inhibition. Likewise, Bat-Hp-CoV and MERS-CoV Nsp1 inhibit translation of mRNAs with non-viral 5’UTRs, whereas reporter mRNAs with their corresponding viral 5’UTRs evaded inhibition (Figure 6B). As a control, Nsp1 proteins with mutations of the key residues in the CTD failed to inhibit protein synthesis (Figure 6B). We next assessed whether viral evasion of Nsp1-mediated translation inhibition occurs in human cells. To this end, we generated isogenic cell lines that stably express *Renilla* luciferase-encoding mRNAs with the 5’UTR from human β-globin, Bat-Hp-CoV, or MERS-CoV upstream of the reporter gene. We compared *Renilla* luciferase (RLuc) signals between the different reporter mRNAs in the presence of WT or mutant Nsp1 proteins. Consistent with our *in vitro* experiments, WT Nsp1 proteins significantly reduced expression of RLuc from the reporter mRNA with a human 5’UTR (Figure 6C, left panel), whereas the corresponding viral mRNA reporters were unaffected in the presence of WT Nsp1 proteins (Figure 6C, right panel).^31^

**Figure 6:**
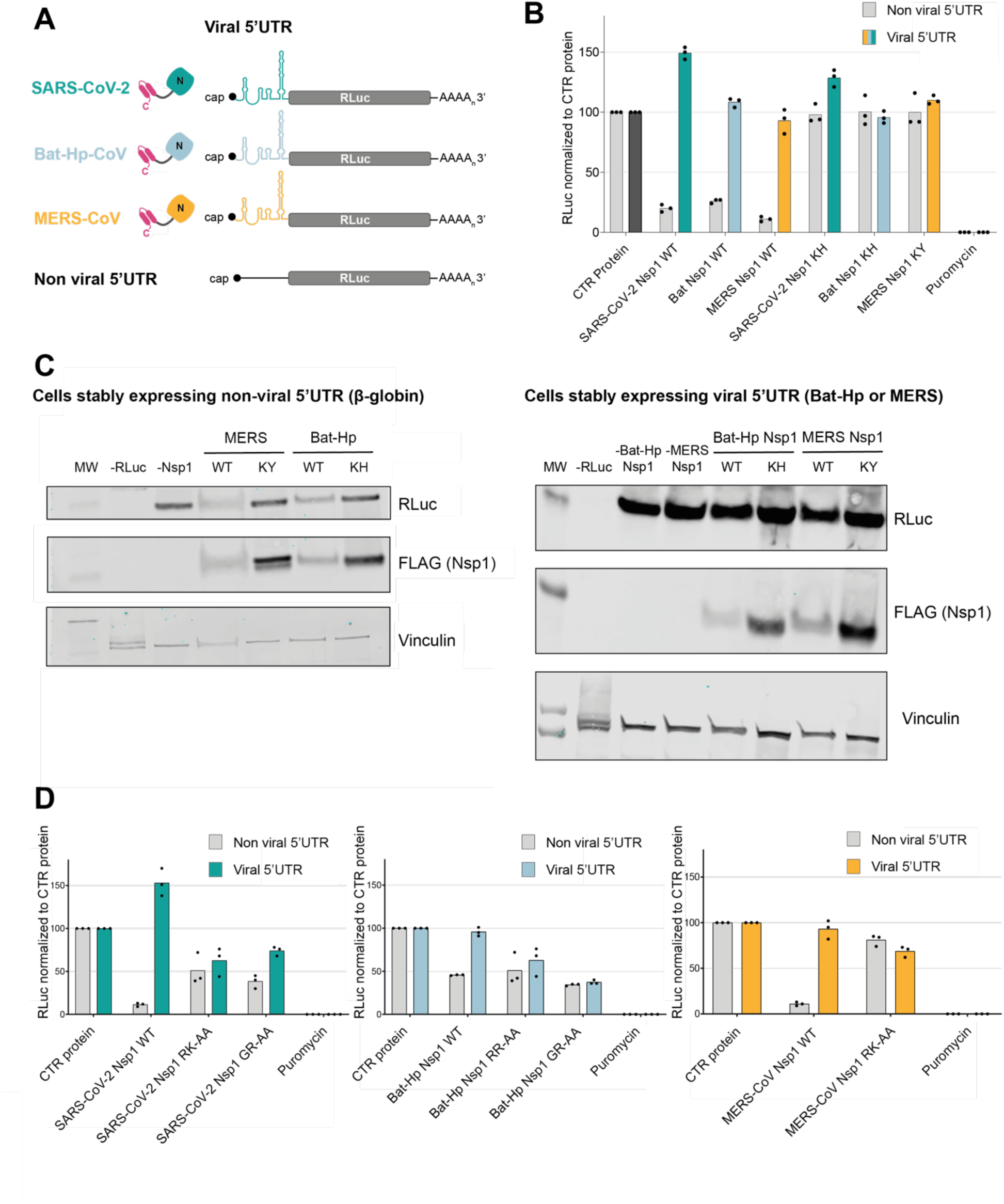
Nsp1 of β-CoVs inhibits host cell translation and protects mRNAs harboring viral 5΄UTRs. **(A)** Schematic of capped and polyadenylated reporter mRNAs coding for *Renilla* Luciferase (RLuc) downstream of viral or non-viral 5΄UTRs. **(B)** RLuc activity measurements of *in vitro* translation reactions containing 300 nM of purified recombinant Nsp1 proteins normalized to the readout of RLuc reporter mRNA in the presence of a control protein. **(C)** Western blot analysis of HEK293 cell lines stably expressing RLuc reporters with 5΄UTRs of human β-globin (hBG), left panel, and Bat-Hp-CoV or MERS-CoV, right panel. hBG-RLuc-expressing cells were transfected with Bat-Hp-CoV or MERS-CoV wild-type FLAG-tagged Nsp1 (WT) or FLAG-tagged Nsp1 with a mutated KH or KY motif for Bat-Hp-CoV (KH) or MERS-CoV (KY), respectively. As a control, two cultures were transfected with an empty plasmid. Cell lysates were prepared, and the Nsp1 expression (anti-FLAG) and the RLuc reporter (anti-RLuc) were determined. **(D)** Relative RLuc activity measurements of *in vitro* translation reactions with 300 nM of purified WT or point mutants of Bat-Hp-CoV, SARS-CoV-2 and MERS-CoV Nsp1, normalized to the reaction at the same concentration of the control protein. Data are presented as mean values of three biological replicates (sets of translation reactions) averaged after three measurements.

Our structural findings predict that substituting the residues within the NTD responsible for binding the decoding center of the 40S subunit will have an opposite effect on the translation of non-viral versus the corresponding viral mRNAs. More specifically, (i) non-viral mRNAs will be translated better in the presence of the mutated Nsp1 relative to the WT protein, since the inhibitory effect of the NTD will be reduced, whereas (ii) viral mRNAs will be translated less efficiently with the mutant Nsp1 proteins relative to WT versions, as previous biochemical results suggest that the same residues are also responsible for interactions with the leader sequence of the viral mRNA^6, 16–18^. Guided by our structure and the sequence conservation of Nsp1, we generated GR-AA substitutions of the decoding center interacting residues for Bat-Hp-CoV (_97_GR_98_) and SARS-CoV-2 (_98_GR_99_) Nsp1 as well as RR-AA/RK-AA mutants in the h18 interacting residues for Bat-Hp-CoV (_123_RR_124_), SARS-CoV-2 (_124_RK_125_) and MERS-CoV (_146_RK_147_) proteins. The effect of all Nsp1 proteins on the translation of reporter genes, preceded by either viral or non-viral 5’UTRs, was assessed using *in vitro* human translation assays. We found a similar trend for Nsp1 from the three β-CoVs. Namely, mutations in either of the two regions of the Nsp1 NTD that contact the 40S subunit alleviated inhibition of the non-viral mRNAs and inhibited translation of the viral mRNAs (Figure 6D), which indicates that the mutant Nsp1 proteins have lost the ability to selectively inhibit translation of host mRNAs.

## Discussion

Identifying common principles for how coronaviruses control translation in host cells is critical to understand and combat coronavirus-related diseases. In this study, we provide new insights into the structure and function of Nsp1 proteins from three different β-CoVs, two of which cause serious human respiratory diseases. We have shown that the binding of the Nsp1 CTD to the mRNA channel of the 40S subunit to inhibit host translation is a conserved mechanism and that the viral mRNAs evade Nsp1-mediated translation inhibition across three divergent β-CoVs. Furthermore, using the Bat-Hp-CoV protein as a model system, we visualized the elusive NTD of Nsp1 bound to the decoding center of the 40S subunit, playing an important role in the selective translation of viral mRNAs (Figure 7).

**Figure 7:**
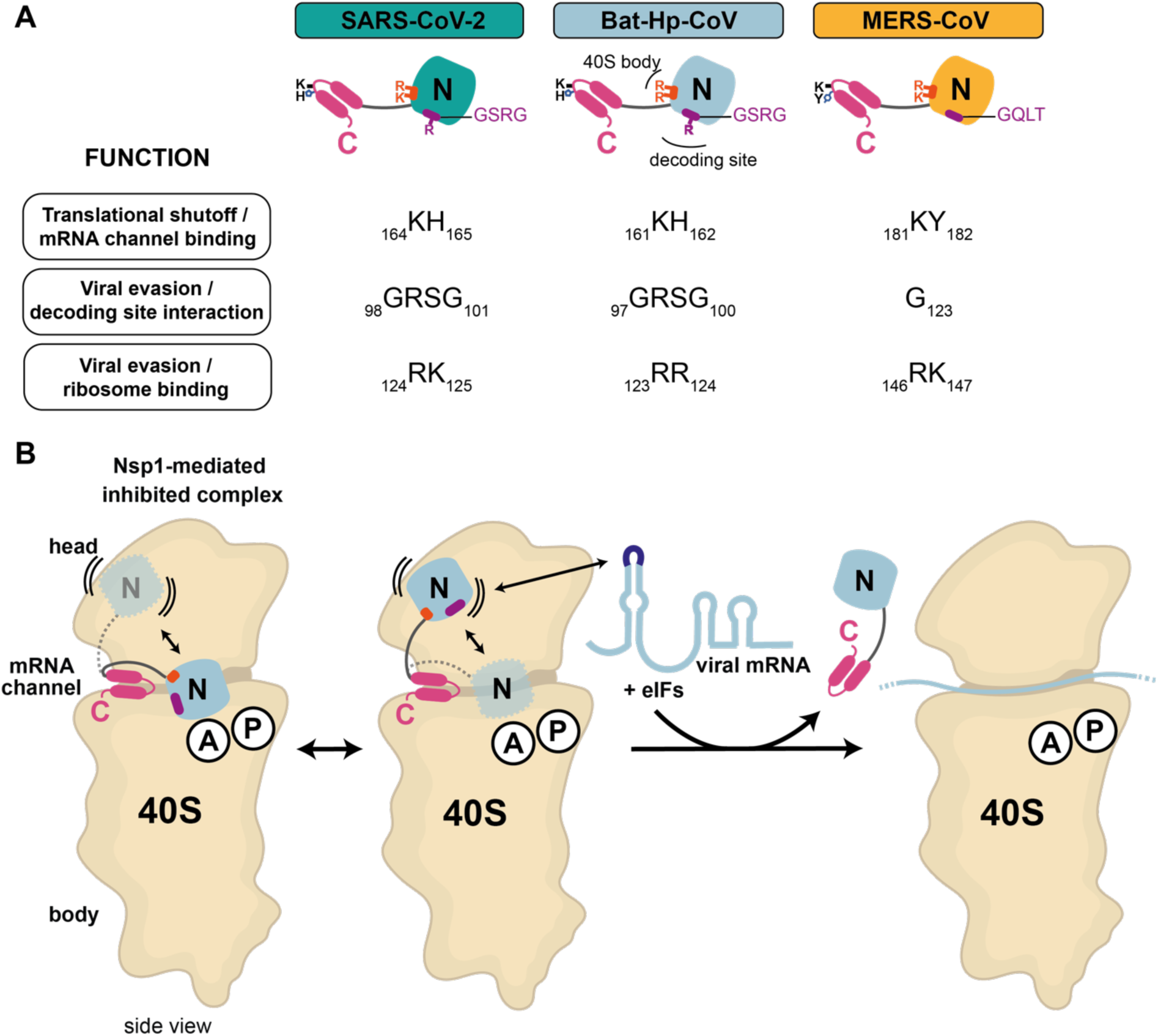
Summary of experimental observations and a mechanistic model for the role of Nsp1 in selective translation of viral mRNAs. **(A)** Schematic overview of SARS-CoV-2, Bat-Hp-CoV and MERS-CoV Nsp1 highlighting amino acids crucial for interacting with the ribosome, inhibiting translation and allowing viral mRNAs to evade the inhibition. **(B)** Nsp1 forms a bi-partite interaction with the 40S subunit, with the CTD binding to the mRNA channel to interfere with mRNA accommodation. The CTD binds with high affinity anchoring Nsp1 to the ribosome, whereas the NTD dynamically interacts with the decoding center, further contributing to translation inhibition by interfering with mRNA accommodation. The 5’UTR of viral mRNAs might hinder docking of the NTD to the decoding center, thereby reducing the inhibition by Nsp1 and leading to preferential translation of viral mRNAs.

Our observation that the NTD of Nsp1 binds to the decoding center of the 40S subunit, the same location where eIF1A would bind in the canonical PICs^32^, allowed us to carry out structure-based experiments aimed to reveal the mechanism of Nsp1 in selective translation shutdown. Although we only visualized the NTD for Bat-Hp-CoV Nsp1, our biochemical data indicate that the binding mode and mechanistic effects on translation apply across all three tested β-CoVs and can be rationalized with our structural data. These findings also explain reported interactions from recent crosslinking-MS studies, in which the Nsp1 linker preferentially localizes on the right side of the ribosomal entry cavity^33^. In agreement with the structural observations, our *in vitro* translation experiments showed that in addition to the CTD, the N-terminal domain also significantly contributes to translation inhibition in all tested viral systems. Furthermore, using structure-guided mutagenesis, we show that the regions of Nsp1 responsible for ribosome interactions are, at the same time, critical for the selective translation of viral mRNAs.

The combination of structural analysis and single-molecule experiments allowed us to evaluate the relative occupancies of Nsp1 domains on the 40S subunit and the dynamics of their interactions. Our results reveal that Nsp1 forms a bi-partite interaction with the 40S subunit. The CTD of Nsp1 binds the 40S mRNA entry channel with high affinity^9, 13–15^ and anchors the protein on the ribosome. Consequently, we observe this interaction for the vast majority of 40S subunits in our imaging conditions. When anchored, the Nsp1 NTD samples multiple conformations. One conformation localizes the NTD to the A-site of the 40S decoding center, as exemplified by the Bat-Hp-CoV protein (∼15% of complexes). The binding of the Nsp1 NTD to the ribosomal A site is mediated in part by the conserved GRSG motif, which directly contacts the decoding bases. In that location, the Nsp1 NTD interferes with eIF1A association and further blocks mRNA accommodation, explaining the effect of the NTD on translation inhibition (Figure 7A). Altogether, our findings are consistent with a model of Nsp1 binding whereby the CTD domain anchors the protein on the 40S subunit with slow dissociation kinetics, and the Nsp1 NTD rapidly samples the 40S decoding center, with only a subset of interactions leading to stable docking. Our inability to detect using structural or single-molecule experiments the NTD of either SARS-CoV-2 or MERS-CoV Nsp1 in the decoding center suggests even faster sampling dynamics and/or less populated residence times for these two viral systems. Despite variable sampling kinetics, an interaction of all three examined Nsp1 proteins with the decoding center is nevertheless supported by our biochemical data when using structure-guided mutants.

Our collective observations suggest a model for how β-CoV RNAs are selectively translated during infection that is consistent with previously published results. To evade Nsp1-mediated inhibition, to which both domains of Nsp1 contribute, special features within the highly structured 5’UTRs of viral mRNAs^20^ may compete for interactions with the NTD with the decoding center of the 40S subunit. Biochemical experiments and sequence analysis indicate a functional interplay between the two. First, the Nsp1 NTDs and the viral mRNAs co-evolved^31^, and consistently, in our experiments we observe translation inhibition evasion when Nsp1 proteins from the three viruses are matched with their corresponding viral mRNAs. Second, mutating either the conserved residues of the Nsp1 NTD revealed here and in previous publications^6, 20^ or the critical nucleotides in stem-loop 1 of the viral 5’UTR, cause the loss of the translation evasion function^20^. Therefore, if viral mRNAs can compete either in *cis* or *trans* with the 40S subunit for interactions with the Nsp1 NTD and occlude its accommodation into the decoding center, the inhibitory potential of Nsp1 will be reduced, leading to preferential translation of viral mRNAs (Figure 7B).

Limitations of this model are that an interaction between the ribosome-bound Nsp1 NTD and the viral mRNA has yet to be observed directly, and that the interaction between the NTD and the decoding center has only been detected for one of the coronaviruses. However, it should be noted that low-affinity interactions between the NTD and viral mRNAs, as well as dynamic interactions between the NTD and the decoding center, are an integral part of the proposed mechanistic explanation: weak interactions with the viral mRNA prevent Nsp1 from being sequestered on viral mRNAs, whereas dynamic interactions with the decoding center allow the viral mRNA access to compete with Nsp1. Regardless, the mechanistic framework provided by our structural, biophysical, and biochemical experiments rationalizes a considerable amount of scientific literature on Nsp1 function. Our work provides a solid foundation to understand how coronaviruses balance suppression of the host immune response with viral protein synthesis and enhances efforts to design anti-viral therapeutics that target Nsp1 activity across *Betacoronaviruses*.

## Acknowledgements

We thank the ETH Scientific center for optical and electron microscopy (ScopeM) and the CryoEM Knowledge hub (CEMK) for technical support. We thank the Functional Genomics Center Zurich (FGCZ) for the help with mass-spectrometry. Thanks to Adrian Bothe, Denis Yudin, Armin Picenoni (all ETH Zurich) and Priyanka Thambythurai (University of Bern) for technical help with initial experiments and Adrian Bothe for carefully reading the manuscript.

This work was supported by grants of NB and OM from the Swiss National Science Foundation (SNSF; 182341,182831, and 204161), the National Center of Competence in Research (NCCR) on RNA and Disease funded by the SNSF (51NF40-182880 and −205601), the ETH Research Grant ETH-23 18-2 to NB, the Multidisciplinary Center of Infectious Diseases from the University of Bern (MCID) to EDK, the Holcim Stiftung Wissen to EDK, and the Forschungsstiftung of the University of Bern to EDK and the U.S. National Institutes of Health (GM011378, GM145306 and AG064690 to J.D.P.; K99GM144678 to C.P.L.).

## Author contributions

CPL and JDP provided the initial Bat-Hp-CoV protein and initiated studies on Bat-Hp-CoV. NB, KS and EDK conceptualized the project. EDK and EB were involved in cloning. KS designed all constructs and expressed all proteins, with the help of TS. KS and IB prepared samples for cryo-EM and froze grids. KS and AS carried out data collection, followed by data processing by KS and IB. KS, TS and IB performed transient transfection and complex purification. EDK designed translation experiments. EDK performed *in vitro* translation reactions, the puromycin incorporation assay and *in vivo* assays, with the help of EB. JZ produced and provided translation-competent lysates. EK analyzed the translation titration experiments. CPL performed the single-molecule experiments and analyses. ML was involved in structure modelling and refinement as well as in figure preparation. NB, KS and EDK coordinated the project. KS and EDK wrote the initial draft. All authors contributed to the final version of the manuscript.

## Declaration of interests

The authors declare no competing interests.

## Tables

**Table S1.** EM data collection and structure refinement statistics.

|  | 40S – Bat-Hp-CoV Nsp1 – eIF1<br>(EMD-XXXXX, PDB XXXX) | 43S PIC – MERS-CoV Nsp1<br>(EMD-ZZZZZ, PDB ZZZZ) |
| --- | --- | --- |
| **Data collection and processing** |  |  |
| Magnification | 81'000x (nominal) |  |
| Voltage (kV) | 300 |  |
| Electron exposure (e<sup>-</sup>/Å<sup>2</sup>) | 60 |  |
| Defocus range (μm) | 0.6-3 |  |
| Pixel size (Å) | 1.065 |  |
| Initial particle images (no.) | 4'054'787 | 1'381'792 |
| Final particle images (no.) | 98'750 | 218'516 |
| Map resolution at FSC=0.143 (Å) | 2.98 | 2.65 |
| **Refinement** |  |  |
| Model resolution at FSC=0.5 (Å) | 2.87 | 2.62 |
| Map sharpening B factor (Å<sup>2</sup>) | -51.0 | -59.3 |
| Model composition |  |  |
| Non-hydrogen atoms | 79'542 | 118'242 |
| Protein residues | 5'189 | 9'771 |
| RNA residues | 1'771 | 1'855 |
| Ligands:<br>Zn<sup>2+</sup>/Mg<sup>2+</sup>/UNX/GTP-Mg<sup>2+</sup>-2xH<sub>2</sub>O | 3/111/117/0 | 4/116/129/1 |
| B factors min/max/mean (Å<sup>2</sup>) |  |  |
| Protein | 37/292/122 | 38/351/198 |
| RNA | 40/576/130 | 35/597/145 |
| Ligand | 40/325/81 | 49/401/219 |
| R.m.s. deviations |  |  |
| Bond lengths (Å) | 0.002 | 0.002 |
| Bond angles (°) | 0.411 | 0.420 |
| **Validation** |  |  |
| MolProbity score | 1.39 | 1.63 |
| Clashscore | 4.75 | 5.95 |
| Poor rotamers (%) | 1.32 | 2.11 |
| Protein |  |  |
| EM Ringer score | 3.3 | 3.00 |
| CaBLAM outliers (%) | 1.5 | 1.4 |
| Ramachandran plot |  |  |
| Favored (%) | 97.75 | 97.93 |
| Allowed (%) | 2.25 | 2.02 |
| Disallowed (%) | 0.0 | 0.05 |
| RNA |  |  |
| Pucker outliers (%) | 0.11 | 0.05 |
| Bond outliers (%) | 0.12 | 0.12 |
| Angle outliers (%) | 0 | 0 |
| Suite outliers (%) | 11.6 | 12.5 |

**Table S2:** Model contents, sequences used for building and modified residues.

| **40S small ribosomal subunit proteins and 18S rRNA** |  |  |  |  |  |  |
| --- | --- | --- | --- | --- | --- | --- |
| **Protein/RNA** | **Alternative (old) name** | **Chain ID <sup>(1)</sup>** | **Full size (residues)** | **Modeled residues <sup>(2)</sup>** | **Uniprot accession code** | **Comments** |
| **uS2** | SA | AA | 295 | 2-217 | P08865 | S2: acetylserine (SAC) |
| **eS1** | S3a | AB | 264 | 2-11, 20-233 | P61247 |  |
| **uS5** | S2 | AC | 293 | 56-278 (59-276) | P15880 |  |
| **uS3** | S3 | AD | 243 | 3-227 | P23396 |  |
| **eS4** | S4 | AE | 263 | 2-263 | P62701 |  |
| **uS7** | S5 | AF | 204 | 13-204 | P46782 |  |
| **eS6** | S6 | AG | 249 | 1-240 | P62753 |  |
| **eS7** | S7 | AH | 194 | 6-194 (6-193) | P62081 |  |
| **eS8** | S8 | AI | 208 | 2-206 | P62241 |  |
| **uS4** | S9 | AJ | 194 | 2-181 | P46781 |  |
| **eS10** | S10 | AK | 165 | 1-97 | P46783 |  |
| **uS17** | S11 | AL | 158 | 2-156 (2-152) | P62280 |  |
| **eS12** | S12 | AM | 132 | 10-132 | P25398 |  |
| **uS15** | S13 | AN | 151 | 2-150 | P62277 |  |
| **uS11** | S14 | AO | 151 | 17-151 | P62263 | D138: isoaspartate (IAS) |
| **uS19** | S15 | AP | 145 | 13-136 (13-143) | P62841 |  |
| **uS9** | S16 | AQ | 146 | 5-146 | P62249 |  |
| **eS17** | S17 | AR | 135 | 2-135 | P08708 |  |
| **uS13** | S18 | AS | 152 | 2-146 | P62269 | S2: acetylserine (SAC) |
| **eS19** | S19 | AT | 145 | 2-145 | P39019 | R67: omega-methylarginine (NMM) |
| **uS10** | S20 | AU | 119 | 17-117 | P60866 |  |
| **eS21** | S21 | AV | 83 | 1-83 | P63220 | M1: acetylmethionine (AME) |
| **uS8** | S15a | AW | 130 | 2-130 | P62244 |  |
| **uS12** | S23 | AX | 143 | 2-142 | P62266 | P62: hydroxyproline (HY3) |
| **eS24** | S24 | AY | 133 | 3-126 | P62847 |  |
| **eS25** | S25 | AZ | 125 | 42-113 | P62851 |  |
| **eS26** | S26 | Aa | 115 | 2-102 | P62854 | bound zinc |
| **eS27** | S27 | Ab | 84 | 2-84 | P42677 |  |
| **eS28** | S28 | Ac | 69 | 4-68 | P62857 |  |
| **uS14** | S29 | Ad | 56 | 2-56 | P62273 | bound zinc |
| **eS30** | S30 | Ae | 133 | 75-133 | NP_001988.1 (GeneBank) | 1-74: ubiquitin |
| **eS31** | S27a | Af | 156 | 78-151 | P62979 | bound zinc, 1-76: ubiquitin |
| **RACK1** | RACK1 | Ag | 315 | 2-315 | P63244 |  |
| **eL41** | L41 | Ah | 25 | 1-25 | P62945 | large subunit protein associated with small subunit |
| **18S rRNA** | - | A2 | 1869 | 1-130, 140-242, 276-706, 724-740, 744-758, 786-1869 | from Taoka *et al.* (RNA 2018) | sequence with modifications according to Taoka *et al.*<sup>34</sup> |

**Table S2 (continued): Model contents, sequences used for building and modified residues.**
| **Additionally built in the Bat-Hp-CoV Nsp1 - eIF1 - 40S complex** |  |  |  |  |  |  |
| --- | --- | --- | --- | --- | --- | --- |
| **Protein** | **Alternative name** | **Chain ID <sup>(1)</sup>** | **Full size (residues)** | **Modeled residues** | **Uniprot accession code** | **Comments** |
| **Bat-Hp-CoV Nsp1** | - | j | 174 | 1-174 | A0A088DIE1 | residues 1-174 of Bat-Hp-betacoronavirus/ Zhejiang2013 polyprotein ORF1ab |
| **eIF1** | EIF1 | p | 113 | 29-113 | P41567 |  |
| **Additionally built in the MERS-CoV Nsp1 - 43S pre-initiation complex** |  |  |  |  |  |  |
| **Protein/RNA** | **Alternative name** | **Chain ID** | **Full size (residues)** | **Modeled residues** | **Uniprot accession code** | **Comments** |
| **eIF1** | EIF1 | lp | 113 | 9-113 | P41567 | rebuilt, N-ter added as poly-Ala |
| **eIF1a** | EIF1AX | lq | 144 | 22-114,126-135 | P47813 | rebuilt |
| **eIF2α** | EIF2S1 | lr | 315 | 4-296 | P05198 | AlphaFold model fitted |
| **eIF2β** | EIF2S2 | ls | 333 | 173-331 | P20042 | bound zinc, AlphaFold model fitted |
| **eIF2γ** | EIF2S3 | lt | 472 | 12-385,393-469 | P41091 | GTP-Mg-2H<sub>2</sub>O bound to active site, AlphaFold model fitted, nucleotide modelled from 4rcy |
| **Met-tRNA<sub>i</sub><sup>Met</sup>** | - | lw | 75 | 1-75 + Met |  | sequence with modifications according to Kowal et al. 2001<sup>35</sup> |
| **eIF3a** | EIF3A | lu | 1382 | 2-588 | Q14152 | AlphaFold model fitted |
| **eIF3c** | EIF3C | ly | 913 | 247-879 | Q99613 | AlphaFold model fitted, linker built as poly-Ala |
| **eIF3d** | EIF3D | lx | 548 | 2-88,157-524 | O15371 | AlphaFold model fitted |
| **eIF3e** | EIF3E | lv | 445 | 1-430 | P60228 | AlphaFold model fitted |
| **eIF3f** | EIF3F | l4 | 357 | 90-357 | O00303 | AlphaFold model fitted |
| **eIF3g** | EIF3G | lo | 320 | 238-314 | O75821 | AlphaFold model fitted |
| **eIF3h** | EIF3H | l8 | 352 | 35-352 | O15372 | AlphaFold model fitted |
| **eIF3k** | EIF3K | l3 | 218 | 2-218 | Q9UBQ5 | 2.1Å crystal structure (1rz4) / human 43S PIC (7a09) as models |
| **eIF3l** | EIF3L | l5 | 564 | 181-552 | Q9Y262 | human 43S PIC (7a09) as model |
| **eIF3m** | EIF3M | l6 | 374 | 11-26, 38-53, 60-372 | Q7L2H7 | human 43S PIC (7a09) as model |
| **MERS-CoV Nsp1** | - | Aj | 191 | 165-193 | K9N7C7 (R1AB_MERS1) | residues 1-193 of MERS polyprotein ORF1ab |

(1) In the Bat-Hp-CoV Nsp1 - eIF1 - 40S complex only a single letter chain ID, corresponding to the 2<sup>nd</sup> letter, is used.
(2) Built ribosomal protein segments that differ in length are additionally given in brackets for the MERS 43S PIC complex.

## STAR Methods

### Cell lines

The lysates for *in vitro* translation experiments were generated from HeLa (ATCC; CCL2) cells that were cultured in DMEM supplemented with 10% FCS, 100 UI/ml penicillin and 100 μg/ml streptomycin (DMEM +/+) at 37°C, 5% CO_2_. For transient transfection of MERS-CoV, 293 c18 (ATCC® CRL-10852™), a human HEK293-EBNA cell line was used. Cells were cultured in EX-CELL 293 serum-free medium at 37°C, 4.5% CO_2_. Both cell lines were tested negative for mycoplasma.

HEK293 Flip-In T-REx cells (Invitrogen #R78007) were cultured as a monolayer in DMEM/F-12 medium (Gibco #32500043) supplemented with 10% FCS, 100 U/ml penicillin and 100 μg/ml streptomycin (37°C, 5% CO2, humidified incubator). SARS-CoV-2, Bat-Hp-CoV, MERS-CoV and human β-globin *Renilla* luciferase constructs were cloned into a modified pcDNA5/FRT/TO (N) plasmid, excluding the tet-operator to allow constitutive expression of the RLuc constructs. Cells stably expressing the aforementioned constructs were established using a pcDNA5/FRT/TO vector-based protocol according to manufacturer’s instructions. The obtained cell lines were tested negative for mycoplasma.

### Cloning, expression and purification of Nsp1 in *E. coli*

Plasmids encoding the SARS-CoV-2 WT Nsp1 sequence and the KH mutant were generated as described^13^. Plasmids encoding Nsp1 wild-type (WT) proteins from MERS-CoV and Bat-Hp-CoV_Zheijang2013 (referred to as Bat-Hp-CoV hereafter) as well as the mutants of SARS-CoV-2 RK-AA (124-125), Bat-Hp-CoV GR-AA (97-98), Bat-Hp-CoV Trx1-linker-CTD and Trx1-SSG-CTD, SARS-CoV-2 Trx1-linker-CTD and Trx1-SSG-CTD as well as MERS-CoV Trx1-linker-CTD and Trx1-SSG-CTD were ordered from GenScript. Mutants of SARS-CoV-2, Bat-Hp-CoV and MERS-CoV were generated by site-directed mutagenesis of the WT using the primers CCCAGGGTTTCACCGCTAGCAGCATACTGAATACCTTCCA to yield SARS-CoV-2 GR-AA (98-99), CCGCGTTCTTGCCACGAGCAGCCAGTTGAATGGTGATGG to yield Bat-Hp-CoV RR-AA (123-124), GATTGCCTTGCGCGGCTGCACCTTTCGGGTCTTCGGTATACAGATC to yield Bat-Hp-CoV KH-AA (161-162), TTCAGCAGGTTTTGCGCAGCCGCGCCCTTCGGGTCCGCTTC to yield MERS-CoV KY-AA (181-182), and GTAGCCACCACGGCCGTATGCAGCCAGCAGGATGTTTTGCTTG to yield MERS-CoV RK-AA (146-147), respectively. The control protein used for normalization in titration assays was a mutated version of Nsp1, in which NTD was replaced by Trx1 and the CTD mutated to KH-AA to inactivate the protein completely. SARS-CoV-2 Trx1-linker-CTD was used as a template with primer GTTCACGGGTCACACCGCTGCTAGCTGCGGTATTCCAATTTTCCTGAAAAT to yield Trx1-linker-CTD KH-AA. Different Nsp1 proteins (WT and mutants from Bat-Hp-CoV, SARS-CoV-2 and MERS-CoV) carrying an N-terminal His_6_-tag followed by a TEV cleavage site were expressed from a pET24a vector. The expression vectors were transformed into *E. coli* BL21-CodonPlus (DE3)-RIPL, and cells were grown in 2xYT medium at 30°C. At an OD_600_ of 0.8, cultures were shifted to 18°C and induced with IPTG added at a final concentration of 0.5 mM. After 16 h, cells were harvested by centrifugation, resuspended in lysis buffer (50 mM HEPES-KOH pH 7.6, 800 mM KCl, 5 mM MgCl_2_, 40 mM imidazole, 10% (w/v) glycerol, 0.5 mM TCEP and protease inhibitors) and lysed using sonification (Sonifer SFX 550). The lysate was cleared by centrifugation for 45 min at 48,000 x g and loaded onto a HisTrap FF 5 ml column (GE Healthcare). Eluted proteins were incubated with TEV protease and dialysed at 4°C overnight using a membrane with a 3.5 kDa MWCO. The His_6_-tag, uncleaved Nsp1 and the His_6_-tagged TEV protease were removed on the HisTrap FF 5 ml column. The sample was further purified via anion exchange chromatography through a 5 ml Q column, or if already sufficiently pure after TEV-cleavage directly *via* size-exclusion chromatography on a HiLoad 16/60 Superdex75 (GE Healthcare), thereby exchanging the buffer of the sample to storage buffer (40 mM HEPES-KOH pH 7.6, 200 mM KCl, 5 mM MgCl_2_, 10% (w/v) glycerol, 1 mM TCEP). Fractions containing Nsp1 were pooled, concentrated in an Amicon Ultra-15 centrifugal filter (10 kDa MWCO), flash-frozen in liquid nitrogen, and stored until further use at −80°C.

### Expression and purification of eIF1 and eIF1A in *E. coli*

Plasmids encoding human eIF1 and eIF1A with an N-terminal His_6_-MBP tag separated by a TEV cleavage site were transformed, expressed, and purified as mentioned^36^. Briefly, the proteins were expressed in *E. coli* BL21-CodonPlus (DE3)-RIPL in 2xYT medium at 18°C. After 16 h, cells were harvested by centrifugation, resuspended in lysis buffer (50 mM HEPES-KOH pH 7.6, 800 mM KCl, 5 mM MgCl_2_, 40 mM imidazole, 10% (w/v) glycerol, 0.5 mM TCEP and protease inhibitors) and lysed using sonification (Sonifer SFX 550). The cleared lysate was loaded onto a HisTrap FF 5 ml column (GE Healthcare) and eluted proteins were incubated with TEV protease, dialysed (3.5 kDa MWCO) at 4°C overnight and run on the HiTrap SP HP 5 ml column to remove cleaved MBP-fusion protein. In case of eIF1, the sample was further purified *via* size-exclusion chromatography on a HiLoad 16/60 Superdex75 (GE Healthcare), buffer exchanging the sample to the storage buffer (40 mM HEPES-KOH pH 7.6, 200 mM KCl, 5 mM MgCl_2_, 10% (w/v) glycerol, 1 mM TCEP). The final protein was flash-frozen in liquid nitrogen and stored until further use at −80°C.

### Transient transfection in HEK293E cells and complex purification

The plasmid carrying the N-terminally 3xFLAG-tagged WT Nsp1 DNA sequence (pcDNA3.1(+) backbone) from MERS-CoV was ordered from GenScript. HEK293E cells were cultured in suspension in EX-CELL 293 serum-free medium at 37°C and 4.5% CO_2_. Approximately 1.8×10^9^ cells were transfected with 2.7 mg of plasmid using PEI MAX (MW 40,000) as transfection reagent (Polysciences, Inc.) in RPMI 1640 medium (Gibco^TM^). After incubation for 1 h, the transfected cells were transferred back into EX-CELL medium supplemented with 3.75 mM valproic acid. After 48h of transient expression, cells were harvested via centrifugation for 10 min at 1,500 x g. After the cell pellet was washed with PBS buffer once, it was flash-frozen in liquid nitrogen and stored at −80°C until further use.

For FLAG-affinity purification, 5 g of cells were lysed in 5 ml of lysis buffer (50 mM HEPES-KOH pH 7.4, 200 mM KOAc, 10 mM Mg(OAc)_2_, 1 mM DTT and 0.5% Triton X-100) using 15 strokes in a dounce homogenizer, followed by incubation for 10 min at 4°C while rotating. To clear the lysate, the sample was centrifuged at 10,000 x g for 15 min. Approximately 0.5 ml of ANTI-FLAG M2 affinity gel were equilibrated in wash buffer (50 mM HEPES-KOH pH 7.4, 200 mM KOAc, 10 mM Mg(OAc)_2_ and 1 mM DTT) and then transferred to the lysate. After incubating the lysate with the beads on a rotary wheel for 1 h at 4°C, the beads were transferred to a Pierce 2 ml centrifuge column (Thermo Fisher Scientific) and washed with 10 column volumes of wash buffer. The sample was eluted with FLAG-peptide (50 mM HEPES-KOH pH 7.4, 200 mM KOAc, 10 mM Mg(OAc)_2_ and 1 mM DTT, 0.2 mg/ml FLAG-peptide). To further purify only the ribosomal complexes, the samples was centrifuged for 2 h at 55,000 rpm (TLA-55 rotor), and the pellet was resuspended in the final cryo-EM buffer (50 mM HEPES-KOH pH 7.4, 100 mM KOAc, 10 mM Mg(OAc)_2_) for 30 min.

### Preparation of human ribosomal subunits

Human ribosomal subunits were purified as described^37^, and final samples were flash-frozen in liquid nitrogen at a concentration of 1 mg/ml (OD_600_ of 10) and stored at −80°C.

### Preparation of viral 5’UTR mRNA and reporter RLuc mRNA

#### *In vitro* transcription plasmids

DNA templates used for *in vitro* transcription reactions were based on a vector with a shortened T7 promoter to reduce the distance between the SL1 loop and the cap, which is crucial for evading translation inhibition by Nsp1. p200-T7mini+G-SARS-CoV2-FL-5΄UTR-6xMS2 was derived after quick change with the mutagenic primer GAC TCA CTA TAG ATT AAA GGT TTA TAC CTT CCC AGG TAA CAA AC using as a template the previously used vector p200-SARS-CoV2-FL-5΄UTR-6xMS2^13^. The modified reporter evaded inhibition in the presence of SARS-CoV-2 Nsp1, in agreement with previous reports^16^. The 5′UTR of Bat-Hp-CoV and MERS-CoV genomic mRNA sequences were subcloned to replace the SARS-CoV-2 5′UTR by fusion PCR using the primers ATG GCT TCC AAG GTG TAC GAC C with CAA TTC ACT GGC CGT CGT TTT ACA A to linearize the vector and ACG GCC AGT GAA TTG TAA TAC GAC TCA CTA TAG TTA AGC TTC GGC TTG with CAC CTT GGA AGC CAT GGT GTC GAG TTG TAG ATG GC to amplify the Bat-Hp-CoV 5′UTR insert cloned in a pUC19 vector as a template, and the SARS-CoV-2 5΄UTR was replaced by the MERS-CoV 5΄UTR genomic mRNA using the pair of primers ACG GCC AGT GAA TTG TAA TAC GAC TCA CTA TAG GAT TTA AGT GAA TAG CTT GGC and CAC CTT GGA AGC CAT GAT GTG CCC CGA ATT GC. The desired plasmids were produced by In-Fusion® HD Cloning (Takara).

#### Eukaryotic expression plasmids

For the transient transfection and expression of 3xFLAG-Nsp1 from Bat-Hp-CoV and MERS-CoV, Nsp1 ORFs were amplified using the corresponding prokaryotic vectors described above as templates with the primers GAC GAT GAC GAC AAA ATG TGC AGC AAG GGC TAC G and CCC TCT AGA CTC GAG TTA GCC GCC GGT CAG GAA CT for Bat-Hp-CoV, CCC TCT AGA CTC GAG TTA ACC GCC AAT CAG T and GAC GAT GAC GAC AAA ATG CAT CAT CAT CAC CAT CAC GAG for MERS-CoV Nsp1 ORFs, and the pcDNA3.1(+)-FLAG vector was amplified with CTC GAG TCT AGA GGG CCC GT and TTT GTC GTC ATC GTC TTT GTA GTC AAT GT. The pairs of PCR fragments were combined to yield the desired plasmids by In-Fusion® HD Cloning (Takara).

#### *In vitro* transcription of reporter mRNAs

Similarly to previously described procedures^13, 38, 39^, plasmids encoding the luciferase reporter mRNA with a T7 promoter in a pCRII vector were linearized using a restriction site located downstream of the poly(A) sequence in a reaction containing 40 ng/μl DNA (4 μg total) and 0.5 U/μl restriction enzyme (*Hind*III-HF, NEB, Cat. No. R3104L) in 1x CutSmart buffer (NEB, Cat. No. B7204S). The digest was monitored by running 200 ng of the DNA on a 1% agarose gel in 1x TAE. The rest of the reaction was purified using the MACHEREY-NAGEL NucleoSpin® Gel and PCR Clean-up kit (Cat. No. 740609.50). The isolated DNA was then quantified by measuring OD_260_.

Using the linearized plasmid as a template (up to 4 μg in total and 40 ng/μl f.c.), the reporter mRNA was *in vitro* transcribed in a reaction containing 1x OPTIZYME transcription buffer (Thermo Fisher Scientific, Cat. No. BP81161), 1 mM of each ribonucleotide (Thermo Fisher Scientific, Cat. No. R0481), 1 U/μl RNase inhibitor (Vazyme, Cat. No. R301-03), 0.001 U/μl pyrophosphatase (Thermo Fisher Scientific, Cat. No. EF0221) and 1 U/μl T7 polymerase (Thermo Fisher Scientific, Cat. No. EP0111). The mixture was incubated at 37 °C for 1 h, and then an equal amount of polymerase was added to boost transcription for another 30 min. Subsequently, the template DNA was degraded by treatment with 0.14 U/μl Turbo DNase (Invitrogen, Cat. No. AM2238) at 37 °C for 30 min. Finally, the newly transcribed reporter mRNA was isolated using the Monarch RNA Cleanup Kit (New England Biolabs, Cat. No. T2040L), eluted with 1 mM sodium citrate pH 6.4 (Gene Link, Cat. No. 40-5014-05) and quantified as mentioned above.

To generate transcripts containing a 7-methylguanylate cap, we used the Vaccinia Capping System from New England Biolabs (Cat. No. M2080S) according to the manufacturer’s instructions with the addition that the reaction mix was supplemented with 1 U/μl RNase inhibitor (Vazyme, Cat. No. R301-03). The resulting capped mRNA was purified, eluted, and quantified as described above, after the transcription reaction.

### Preparation of HeLa translation-competent lysates

Based on a previously established protocol^39^, lysates were prepared from HeLa S3 cell cultures grown to a cell density ranging from 0.8×10^6^ to 1×10^6^ cells/ml. Cells were pelleted (200 x *g*, 4°C for 5 min), washed twice with cold 1x PBS pH 7.4 and resuspended in a translation-competent (lysis) buffer (33.78 mM HEPES pH 7.3, 63.06 mM KOAc, 0.68 mM MgCl_2_, 54.05 mM KCl, 13.51 mM creatine phosphate, 229.73 ng/ml creatine kinase and 1x protease inhibitor cocktail (Bimake, Cat. No. B14002)) at a final concentration of 2×10^8^ cells/ml in 2 ml screw cap microtubes (Sarstedt, Cat. No. 72.693). Cells were lysed by dual centrifugation at 500 rpm at −5°C for 4 min using ZentriMix 380 R system (Hettich AG) with a 3206 rotor with 3209 adapters. The lysate was centrifuged at 13,000 x *g*, 4°C for 10 min and the supernatant (the lysate) was aliquoted, snap-frozen and stored at −80 °C. The lysate could be thawed and refrozen multiple times for subsequent applications with only a minor loss of translation efficiency.

### *In vitro* translation assays

*In vitro* translation reactions were performed similarly as described before^13^. Briefly, for SARS-CoV-2 Nsp1 titration reactions, 400 μl of recombinant proteins were dialyzed overnight in Buffer A (30 mM NaCl, 5 mM Hepes pH 7.3) at 4 °C using Slide-A-Lyzer MINI Dialysis devices with a 3.5 kDa MWCO (Thermo Scientific, 88400), and the protein concentration was calculated using Nanodrop. To assess possible contaminants that may affect the translation efficiency, a control protein (Trx1-SARS-CoV-2-CTD-KH) was purified in parallel and applied at an equal concentration of Nsp1 proteins in every assay. For titration experiments, the highest concentration was chosen (Figure S8). For MERS-CoV and Bat-Hp-CoV Nsp1 constructs as well as the Trx1-Nsp1-CTD-KH (control protein), the proteins were diluted with Buffer A to the desired concentrations. In parallel, equal volumes of recombinant protein storage buffer were either dialyzed or diluted to the same final concentration as contained in the diluted proteins and used as negative control (0 μM condition). HeLa S3 lysates prepared in hypotonic lysis buffer were used at a concentration of 8.88 x10^7^ cell equivalents/ml. To perform *in vitro* translation, the lysates were supplemented with 1 u/μl RNase inhibitor (Vazyme, Cat. No. R301-03). Before adding reporter mRNAs, the mixtures were incubated at 33°C for 5 min. *In vitro* transcribed and capped mRNAs with non-viral 5΄UTRs were incubated for 5 min at 65°C and cooled on ice. Reporters with viral 5΄UTRs were incubated for 5 min at 65°C, followed by 15 min at room temperature, and were then placed on ice until usage. Reporter mRNAs were added to the translation reactions at a final concentration of 5 fmols/μl, the translation reaction was performed at 33°C for 50 min and was terminated by transferring the samples on ice. To monitor the protein synthesis output by the luciferase assay, 12.5 μl of each translation reaction was mixed with 50 μl 1x *Renilla*-Glo substrate in *Renilla*-Glo assay buffer (Promega, Cat. No. E2720) on a white-bottom 96-well plate (Greiner, Cat. No. 655073). The luminescence signal was measured three times using the TECAN infinite M1000 Pro plate reader according to the manufacturer’s guidelines and plotted on GraphPad (Version 8.4.2). Data processing for the titration assays was done with the statistical software R 4.2.2.

### Transient expression of FLAG-Nsp1 in HeLa cells, puromycin incorporation assay and western blot analysis

HEK293 cells were cultured in DMEM supplemented with 10% FCS and with 100 UI/ml penicillin and 100 μg/ml streptomycin (DMEM+/+) at 37 °C, 5% CO_2_. To compare expression levels of different FLAG-Nsp1 protein variants (WT/KY), ∼4 × 10^5^ cells were transfected with 400 ng of each plasmid using 7.5 μl μg^−1^ DNA Dogtor transfection reagent (Oz Biosciences) in Opti-MEM (Thermo Fisher Scientific). As a control, an equal amount of empty vector was added to the reaction. After 24 h the samples were collected by trypsinization, counted and lysed in RIPA buffer (50 mM Tris-HCl pH 8.0, 150 mM NaCl, 1% NP-40, 0.5% sodium deoxycholate, 1% SDS). The whole-cell extract was then isolated by centrifugation at 13,000 x *g* and 4°C.

For the puromycin incorporation assay, ∼1.25 × 10^5^ cells were transfected with 500 ng of the WT FLAG-Nsp1 plasmid (or empty vector) and 100 ng of the FLAG-Nsp1 mutant plasmids (using Dogtor and Opti-MEM as above). After one day, the transfection was repeated with double the amount of plasmid DNA and the cells were incubated for 24 h again. Forty minutes before the end of this incubation time the medium was changed to DMEM+/+ containing 10 μg/ml puromycin (Santa Cruz Biotechnology) for 10 min to label newly synthesized proteins. Subsequently, the cells were washed with 1x PBS, and the cells were recovered in the original medium for 30 min. Ultimately, the cells were harvested in 1x PBS. In addition, a positive control condition for translation inhibition was included (empty vector transfected), in which the medium was supplemented with CHX (Focus Biomolecules) to 100 μg/ml within the last 2 h of the experiment (apart from the puromycin pulse).

All western blot samples were dissolved in 1.5x LDS loading buffer (Invitrogen, NP0008) containing 50 mM DTT and were incubated at 75°C for 5 min. The samples were separated on a mPAGE 4-12% Bis-Tris Precast Gel (Millipore, Cat. No. MP41G15) in MES buffer (50 mM MES, 50 mM Tris base, 0.1% SDS, 1 mM EDTA, pH 7.3) at 150 Volts. The gel was transferred onto a nitrocellulose membrane with the Trans-Blot Turbo Transfer system using a preassembled Transfer Pack Midi (Cat. No. 1704159, Bio Rad). The membrane was blocked with 5% BSA in TBS-T (0.01% NaN_3_) for 30 min. Mouse anti-Vinculin (1:2000, Santa Cruz, cat. no. sc-73614), Rabbit anti-*Renilla* luciferase (Thermo Fisher Scientific, Cat. No. PA5-32,210, 1:500) and Mouse anti-Flag (1:2500, Sigma Aldrich, cat. no. F1804) primary antibodies were added and incubated overnight at 4°C. The membrane was rinsed with TBS-tween (0.1%) three times for five minutes. The membrane was incubated with Donkey anti-rabbit 800CW (LI-COR, Cat. No. 926-32212, 1:10’000), Donkey anti-mouse 700CW (LI-COR, Cat. No. 926-68022, 1:10’000), followed by rinsing as before. The signals were visualized using an Odyssey Infrared Image System at 700 and 800 nm.

### Single-molecule spectroscopy

#### TIRFm imaging

Data were collected on a home-built, prism-based instrument. All data were collected at room temperature at 10 frames per second with the EM gain set to 650. Quartz slides were coated with a methoxy-PEG and biotin-PEG mixture using a protocol derived from Ha and colleagues^40^. Fluorescence emission data were collected in both the Cy3 (donor) and Cy5 (acceptor) channels following excitation of the Cy3 donor dye with the 532 nm laser. Data reported in Figure 5B, 5C (SARS-CoV-2 only), and Figure S7A,B,C were obtained using TIRFm.

#### ZMW imaging

Equilibrium and real-time imaging in zero-mode waveguides (ZMWs) were conducted using a modified Pacific Biosciences RSII DNA sequencer and the Maggie software (v. 2.3.0.3.154799), which were described previously^41^. Experiments were performed at 20 °C or 30 °C (as indicated) using a 532 nm excitation laser at 0.32 µW/µm^2^, which directly excited Cy3 and Cy3.5 dyes. Cy5 dye was excited via FRET as indicated. Fluorescence emission data (Cy3, Cy3.5, and Cy5) were collected at 10 frames per second for 600 sec. ZMW chips were purchased from Pacific Biosciences, which were passivated by reaction with polyvinylphosphonic acid to form a covalent Al-polyphosphonate coating^42^. Unless noted above in the TIRFm section, all other single-molecule experiments were performed using ZMW imaging.

#### Surface preparation

Prior to imaging, the quartz slide surface or ZMW surfaces were washed with 0.2% Tween-20 and TP50 buffer (50 mM Tris-OAc pH 7.5, 100 mM KCl). Washed surfaces were coated with neutravidin by a 5 min incubation with 1 µM neutravidin diluted in TP50 buffer supplemented with 0.7 mg/ml UltraPure BSA and 1.3 µM of pre-annealed DNA blocking oligos (CGTTTACACGTGGGGTCCCAAGCACGCGGCTACTAGATCACGGCTCAGCT) and (AGCTGAGCCGTGATCTAGTAGCCGCGTGCTTGGGACCCCACGTGTAAACG). The imaging surfaces then were washed with TP50 buffer at least four times.

#### Reaction and Imaging buffers

For all single-molecule assays, the ‘reaction buffer’ was: 20 mM HEPES-KOH, pH 7.3, 100 mM KOAc, and 2 mM Mg(OAc)_2_. The ‘imaging buffer’ was the reaction buffer supplemented with casein (62.5 µg/ml) and an oxygen scavenging system^43^: 2 mM TSY, 2 mM protocatechuic acid (PCA), and 0.06 U/µL protocatechuate-3,4-dioxygenase (PCD).

#### Labeled 40S subunits and Nsp1 proteins

Human 40S subunits labeled with Cy3 dye on the N-terminus of uS19 were purified and fluorescently labeled as described^15^. SARS-CoV-2 and Bat-Hp-CoV Nsp1 proteins were purified and fluorescently labeled with Cy5 dye also as described^15^.

#### RNA identity and preparation

Since we were unable to biotinylate and fluorescently-label 40S subunits simultaneously, we relied on an RNA that directly binds the 40S subunit outside the mRNA entry channel and decoding center regions to tether Cy3-labeled 40S subunits to imaging surfaces. The RNA was based on the hepatitis C virus (HCV) internal ribosome entry site (IRES) and was described and used previously by our group^15^. RNAs lacked nucleotides downstream of the annotated translation start site (AUG-3’; the described +0 reporters).

#### Complex formation for equilibrium TIRFm and ZMW experiments

As the RNA, we used our described version that lacked domain II of the HCV IRES (Δ-dII)^15^, which directly manipulates the 40S head conformation. With this Δ-dII RNA, the 40S head adopts the closed conformation, which brings the uS19 and Nsp1 labeling sites to closer proximity. However, Nsp1 binds slowly to the 40S subunit in this conformation^15^. We therefore assembled the complex step wise to maximize Nsp1 occupancy on the 40S subunit. First, 75 nM Cy3-40S subunits, 200 nM Nsp1-Cy5 (SARS-CoV-2 or Bat-Hp-CoV), and 500 nM eIF1 were incubated for 10 min at 37 °C in reaction buffer. Second, 20 nM 5’-biotinylated RNA was added to the mixture and incubated for an additional 5 min. The resulting complex was then tethered to TIRFm or ZMW imaging surfaces, washed once with imaging buffer, and imaged as reported above. During imaging, the imaging buffer was supplemented with 10-20 nM Cy5-Nsp1 and 200 nM eIF1 to maintain their occupancy on tethered 40S subunits.

#### Real time competition assays

To examine Nsp1 and eIF1A association with tethered 40S subunits in real time, we again leveraged the HCV IRES. Since Nsp1 binds slowly to 40S subunits bound to the 1′-dII version of the IRES, we used the WT version that contained domain II (dII), which promotes rapid Nsp1 association with 40S subunits via induction of the open head conformation^15^. To form the RNA-40S complex, 75 nM Cy3-labeled 40S subunits were incubated with 20 nM RNA in reaction buffer for 20 min at 37 °C. The resulting complex was tethered to the ZMW surface for 5 min and washed once with imaging buffer. At the onset of data acquisition, 25 nM of Cy5-eIF1A and 25 nM of Cy3.5-Nsp1 (SARS-CoV-2 or Bat-Hp-CoV) were added to the tethered complex.

### Single-molecule data analyses

Experimental movies that captured fluorescence intensities over time were processed and analyzed using MATLAB R2018-b as described previously^15, 41, 44, 45^. To determine *E_FRET_*, fluorescence data from individual TIRFm spots or ZMWs with the desired fluorescence signals (e.g., 40S-Nsp1 FRET) were background-corrected using SPARTAN 3.7.0^46^, with single molecules indicated by single-step photobleach events of the donor fluorophore. FRET on and off states were assigned automatically using vbFRET (version june10)^47^, which were visually inspected and manually corrected as needed. An *E_FRET_* threshold of 0.1 was used in all experiments. *E_FRET_* was defined as:

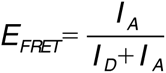

Where *I_D_* and *I_A_* represent fluorescence intensities of the donor and acceptor fluorophores. *E_FRET_* values observed across all events and molecules were binned (50 bins, −0.2 to 1.2) and fit to gaussian functions to determine the mean and standard deviations. To determine association and dissociation kinetics, binding events of individual components (e.g., Nsp1, eIF1A) were assigned manually based on the appearance and disappearance of the respective fluorescence signals. eIF1A association times and binding lifetimes were defined as the time elapsed from the previous binding event until appearance of the next binding event (association), and the duration of each binding event (lifetime). The observed association times and lifetimes were used to calculate cumulative probability functions of the observed data (cdfcalc, MATLAB), which were fit to single- or double-exponential functions in MATLAB (cftool, non-linear least squares method). Derived association rates, lifetimes, and the number of molecules examined are reported in the relevant figures and legends. The exponential function was defined as:

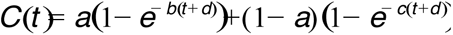

where *t* is time (in seconds), *b* and *c* are rates, and *d* is an adjustment factor. If the analysis yielded a phase that represented less than 10% of the population, only a single phase was used to derive the respective rate and reported (i.e. single-exponential function).

#### Statistical analyses

To calculate errors for the efficiency of a given binding event, bootstrap analyses (n = 10,000) were performed to calculate 95% confidence intervals (C.I.) for the observed proportions using R (4.1.2) and RStudio (Ghost Orchid release) (mosaic library, rflip approach). Reported errors for derived rates represent 95% C.I. yielded from fits to linear, single-exponential, or double-exponential functions, as indicated.

### Cryo-EM sample preparation and data collection

Quantifoil R2/2 holey carbon copper grids (Quantifoil Micro Tool) were prepared by first applying an additional thin layer of continuous carbon and then glow-discharging them for 15 sec at 15 mA using an easiGlow Discharge cleaning system (PELCO). For the *in vitro* binding experiment, purified Bat-Hp-CoV Nsp1 and human eIF1 were mixed with human 40S in molar ratios of 2.5 : 2.5 : 1 and adjusting the 40S to a concentration of 80 nM. For the FLAG-purified MERS-CoV Nsp1 ribosomal complexes, the final sample concentration was adjusted to 100 nM. For both complexes, 4 µL of the sample was applied to the grids, which were then blotted for approximately 1-2 s and immediately plunged into a 1:2 ethane:propane (Carbagas) mixture at liquid nitrogen temperature using a Vitrobot (Thermo Fisher Scientific). The Vitrobot chamber was kept at 4°C and 100% humidity during the whole procedure.

For each sample, one grid was selected for data collection using a Titan Krios cryo-transmission electron microscope (Thermo Fisher Scientific) operating at 300 kV and equipped with a K3 camera (Gatan), which was run in counting and super-resolution mode, mounted to a GIF Quantum LS operated with an energy filter slit width of 20 eV. The K3 datasets were collected at a nominal magnification of 81,000x (physical pixel size of 1.065 Å/pixel). For counting mode, illumination conditions were adjusted to an exposure rate of 24 e^-^/pixel/sec. Micrographs were recorded as movie stacks at an electron dose of ∼60 e^-^/Å^2^ applied over 40 frames. For both datasets, the defocus was varied from approximately −1 to - 3 μm.

### Cryo-EM data processing

The stacks of frames were first aligned to correct for motion during exposure, dose-weighted, gain-corrected, and their CTF was estimated using cryoSPARC Live^48^.

Micrographs (8,932 for MERS-CoV 43S PIC, 12,607 for the *in vitro* Bat-Hp-CoV Nsp1 – eIF1 – 40S complex) were carefully inspected based on CTF estimations, overall resolution as well as for drift and ice quality. Particle images of ribosomes were picked (1,381,792 for MERS-CoV 43S PIC, 4,054,787 for the *in vitro* Bat-Hp-CoV Nsp1 – eIF1 – 40S complex) in cryoSPARC^48^ using the template picker option with EMD-11320 and EMD-11609 as inputs for the MERS-CoV 43S PIC and EMD-11320 as input for the Bat-Hp-CoV Nsp1 complex. The picked particle images were then subjected to a reference-free 2D classification in cryoSPARC, and the particles were selected from the 2D class-averages (888,759 for MERS-CoV 43S PIC, 1,276,183 for the *in vitro* Bat-Hp-CoV Nsp1 – eIF1 – 40S complex).

For the MERS-CoV 43S PIC dataset, the particles were then classified in 3D using human 40S and 80S – SARS-CoV-2 Nsp1 complexes (EMD-11320 and EMD-11609) as inputs that were low-pass filtered to 20 Å to select for good 40S and 80S classes (Figure S1). Within the refined classes, two main states were identified and further processed: a translation inactive 80S complex with bound eEF2 and SERBP1^13, 49, 50^ and a partially resolved 43S PIC. Since we were interested in Nsp1-bound complexes, only the 43S PIC was further analyzed by performing a global 3D variability analysis in cryoSPARC^51^. Two main classes were sorted, one including the 43S PIC including strong density for eIF3 (218,516) and one without eIF3 density (368,744). The 43S PIC with all the initiation factors bound was then un-binned to full pixel size (1.065 Å/pixel) and refined to a final resolution of 2.65 Å.

For the Bat-Hp-CoV Nsp1 complex dataset, the particles were also classified in 3D using the human 40S – SARS-CoV-2 Nsp1 complex (EMD-11320) as input low-pass filtered to 20 Å (Figure S5). High-quality classes were merged and subjected to global 3D variability analysis^51^ yielding three distinct classes of the 40S ribosome: two Nsp1 + eIF1 bound states, one with open 40S head and one in a slightly rotated head state as well as one class of 40S without any factors present. For both head rotation states, a mask around Nsp1 NTD and eIF1 was created in UCSF Chimera^52^ and another round of 3D variability analysis was performed, yielding to states with partial densities of factors. The final classes were un-binned to full pixel size and refined to 2.98 Å for the open head state (98,750 particles) and to 3.16 Å for the rotated head state (9,593 particles).

### Model building and refinement

For initial interpretation of the human 40S – Bat-Hp-CoV Nsp1 cryo-EM map, individual structures of the human small ribosomal subunit head (6zol), as well as the 40S body in complex with the Nsp1 C-terminus of SARS-CoV-2 (6zok)^13^, were docked as rigid bodies using Chimera^52^, combined and manually adjusted in COOT^53^. For initial docking of the Nsp1 NTD, a model generated by Alphafold2^54^ was used, while the SARS-CoV-2 Nsp1 CTD was mutated to the Bat-Hp-CoV_Zhejiang2013 sequence. The 18S rRNA modifications were built according to the quantitative mass spectrometry data obtained by Taoka et al.^34^ taking the closely related rabbit 18S high resolution structure (7o7y)^55^ as a guide. While a few rRNA modifications were not visible in the maps due to low occupancy or flexibility and were omitted, several N-terminal and internal protein modifications were clearly recognizable and were also included in the 40S model (Table S1).

The ribosomal part of the human 43S PIC – MERS-CoV Nsp1 complex was built as described above. For interpretation of the additionally bound initiation factors (Table S2), models generated by Alphafold2^54^ were docked and manually adjusted, except for the peripheral, less-well resolved eIF3K, eIF3L and eIF3M subunits, which were transplanted from a published human 43S pre-initiation complex (7a09)^56^. Few protein linker segments or terminal extensions with weak density were modelled as unassigned UNK residues (Table S2). For the bound human Met-tRNAi(Met), a published modification pattern was built^35^. The GTP nucleotide bound to the eIF2 gamma subunit was modelled according to a high-resolution archaeal structure (4rcy)^57^.

Both structures were refined using phenix.real_space_refine (version 1.20.1)^58^. For this, the coordinates were subjected to multiple refinement cycles including protein rotamer and Ramachandran restraints. The resulting models were validated and improved using Molprobity^59^ and the tools implemented in COOT^53^. Discrepancies between the maps and models were detected and corrected with the help of real space difference maps. Only ions known to be hexa-coordinated magnesiums from comparison with the high-resolution rabbit model (7o7y)^55^ were modelled as such, while the remaining ion densities were interpreted as unknown ions (UNXs). The final overall map-to-model fit was evaluated using the real space correlation coefficients (Table S2) and the model vs. map FSCs at a value of 0.5, with resolutions coinciding well with those obtained from the half maps at the FSC=0.143 criterion (Figure S1 and S5).

## Data availability

Cryo-EM maps and atomic models have been deposited in the Electron Microscopy Data Bank (EMDB) and wwPDB, respectively, with the following accession codes: EMD-XXXXX and PDB XXXX for the Bat-Hp-CoV Nsp1 – IF1 – 40S complex and EMD-ZZZZZ and PDB ZZZZ for the MERS-CoV Nsp1 – 43S PIC.

## Supplemental information

**Figure S1:**
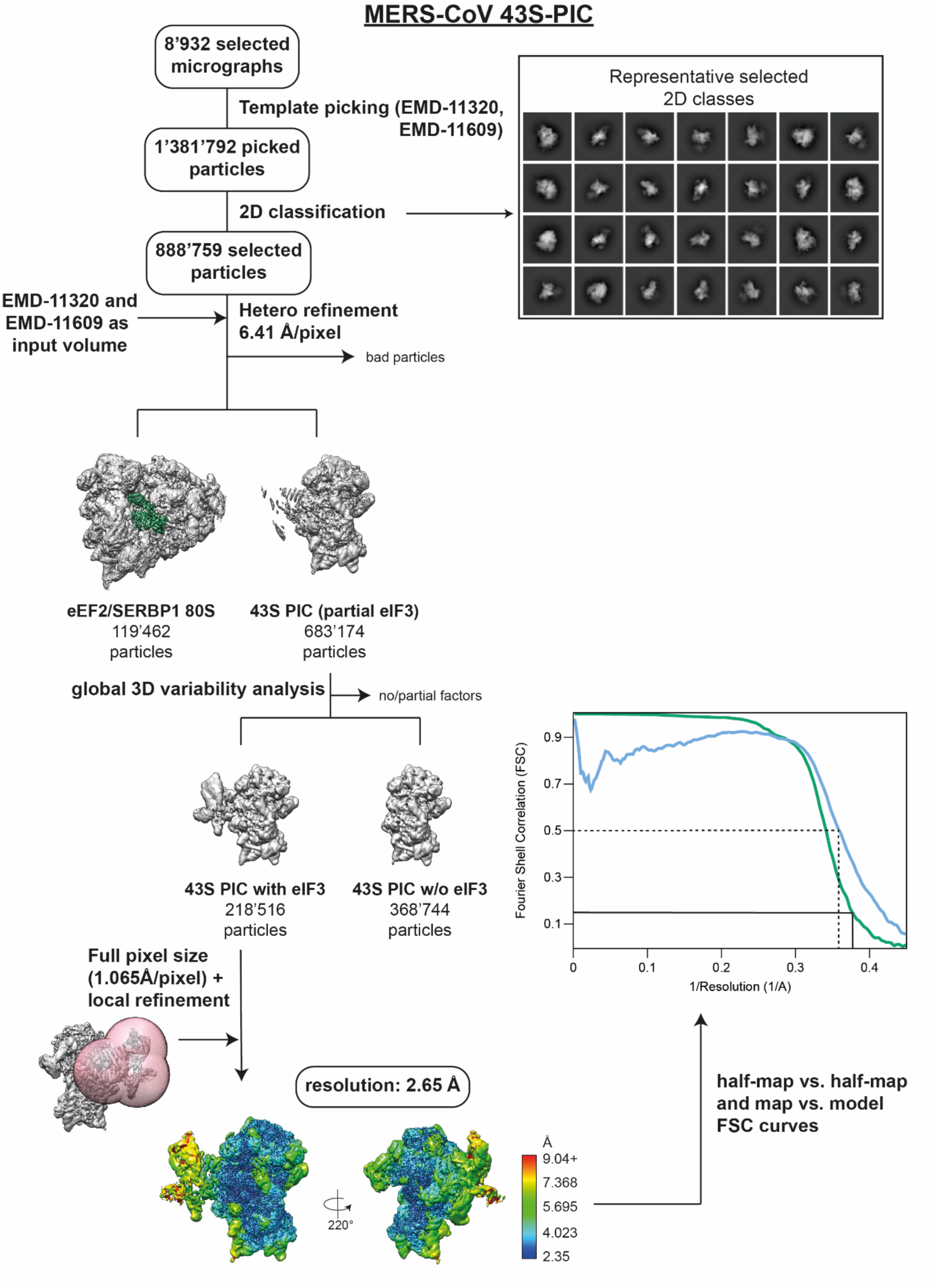
Scheme for the processing of FLAG-MERS-CoV Nsp1 sample purified from HEK293E cell lysate. Local resolution estimates are plotted as heat map on the final volumes accompanied by the color scheme. The half map vs. half map and map vs. model FSC curves are shown for the refinement.

**Figure S2:**
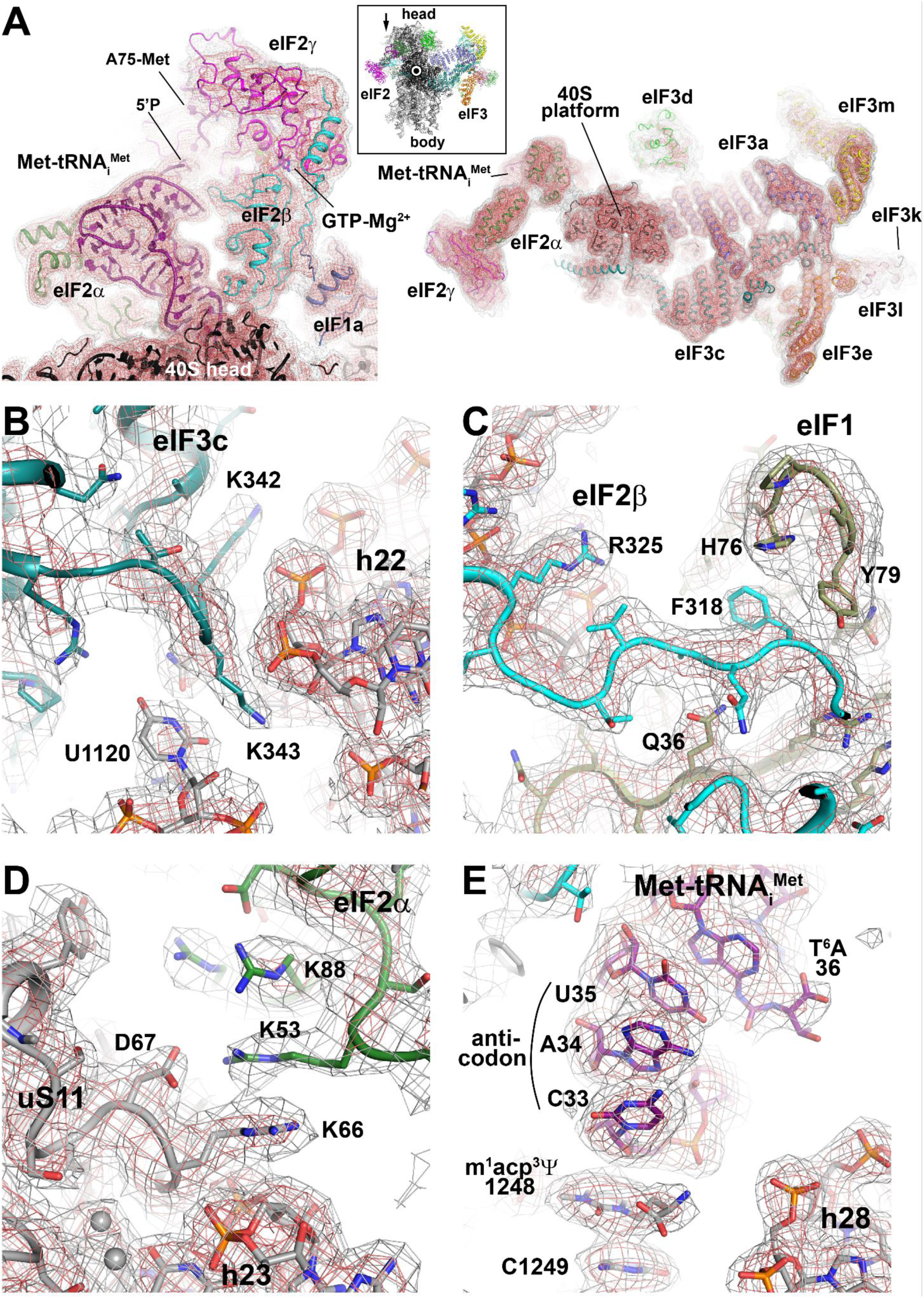
EM densities of the MERS-CoV Nsp1-bound human 43S pre-initiation complex. **(A)** The maps filtered to 5Å reveal the overall fit of the ternary IF2 complex (left) and the IF3 subunits (right). For orientation, the inset indicates the view directions (from top as an arrow or along the view axis as a circle). **(B)** – **(E)** Representative examples of the EM density close to the 40S, where it reaches near-atomic resolution. 40S rRNA and ribosomal proteins are colored in grey.

**Figure S3:**
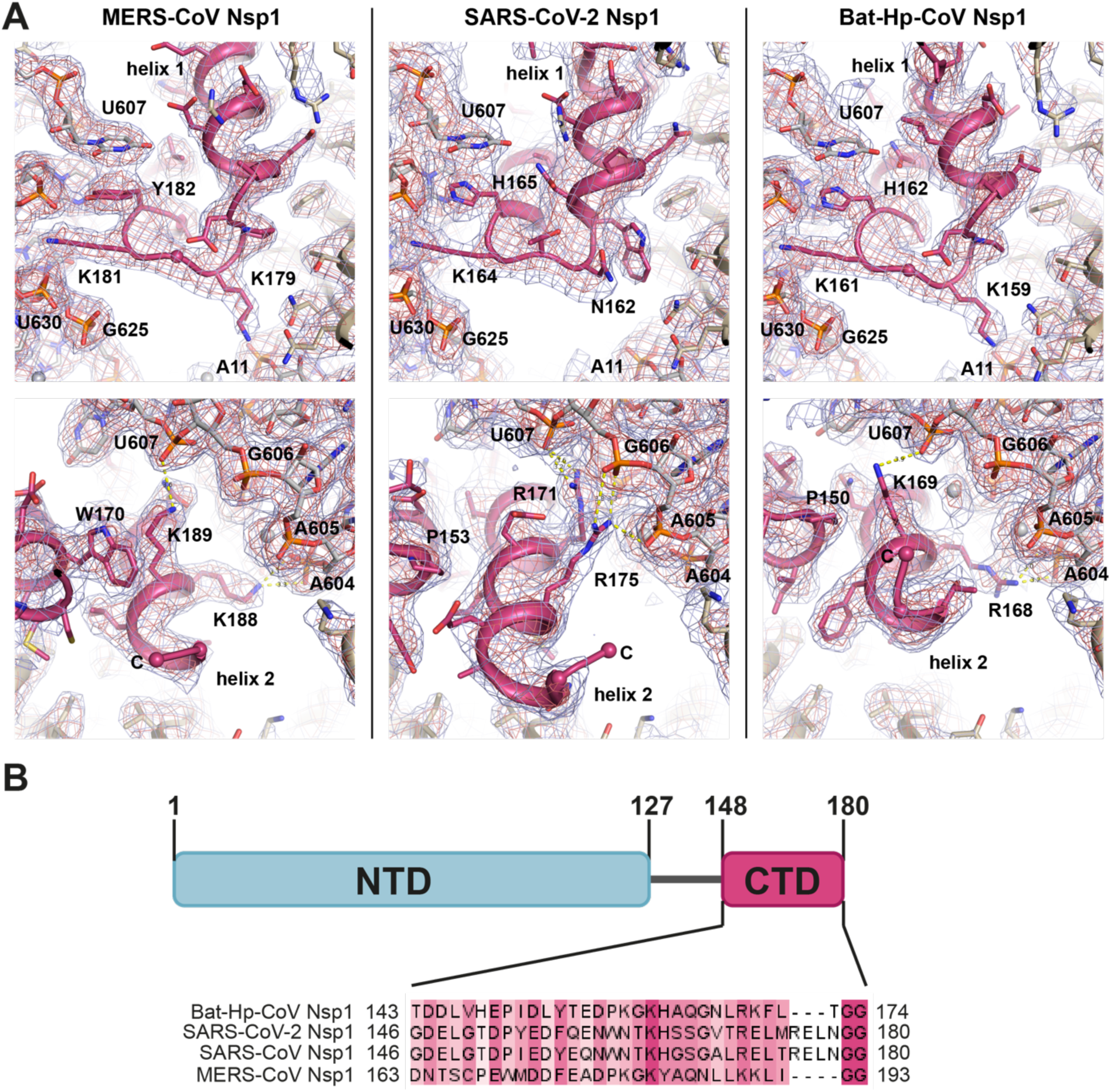
Molecular interactions of the C-terminal domains of MERS-CoV, SARS-CoV-2 and Bat-Hp-CoV Nsp1 are conserved. **(A)** Detailed views of the specific interactions of Nsp1 binding (magenta) to uS5 and 18S rRNA. The experimental densities are shown in red and dark blue mesh contoured at 7σ and 11σ, respectively. **(B)** Clustal Omega sequence alignment of Nsp1 from Bat-Hp-CoV, SARS-CoV-2, SARS-CoV, and MERS-CoV. Colored according to conservation in Jalview with an overall conservation threshold of 20^30^.

**Figure S4:**
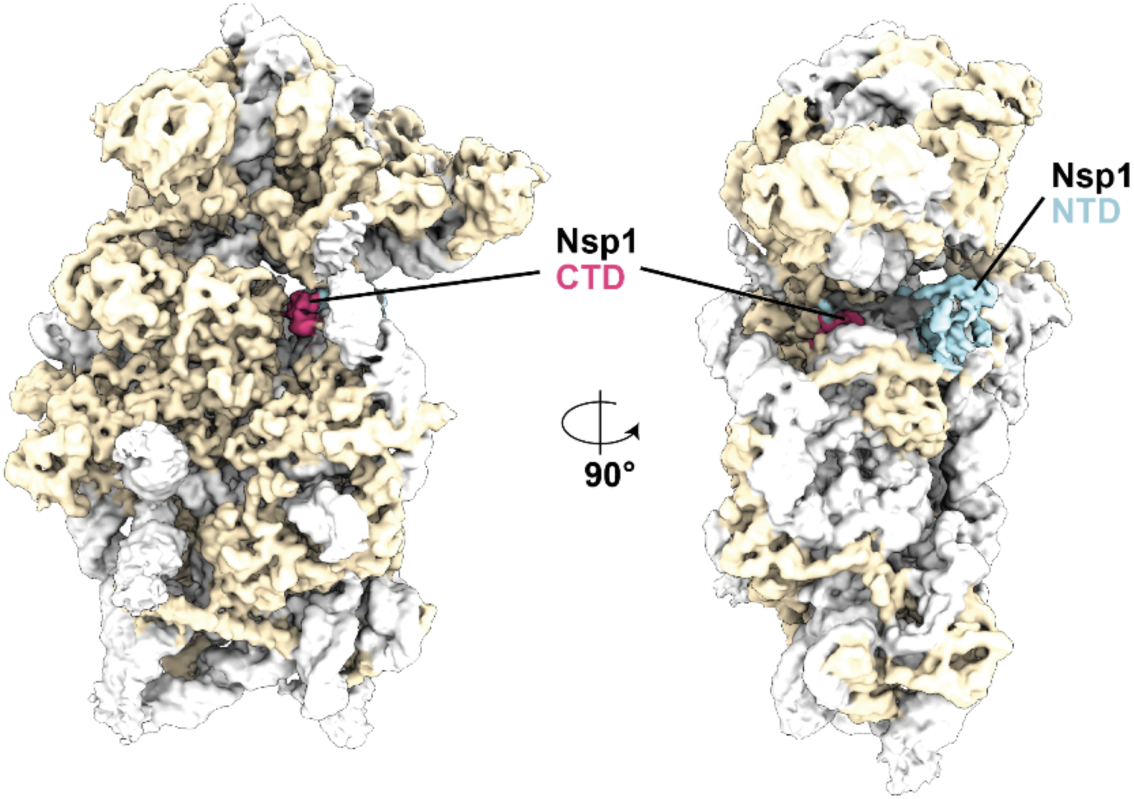
Binding interface of the CTD and NTD of Bat-Hp-CoV Nsp1 in complex with the 40S subunit without the addition of eIF1. Views of Bat-Hp-CoV Nsp1 (CTD in magenta and NTD in light blue) binding to the small ribosomal subunit (rRNA in light grey, ribosomal proteins in beige). The map is filtered to local resolution.

**Figure S5:**
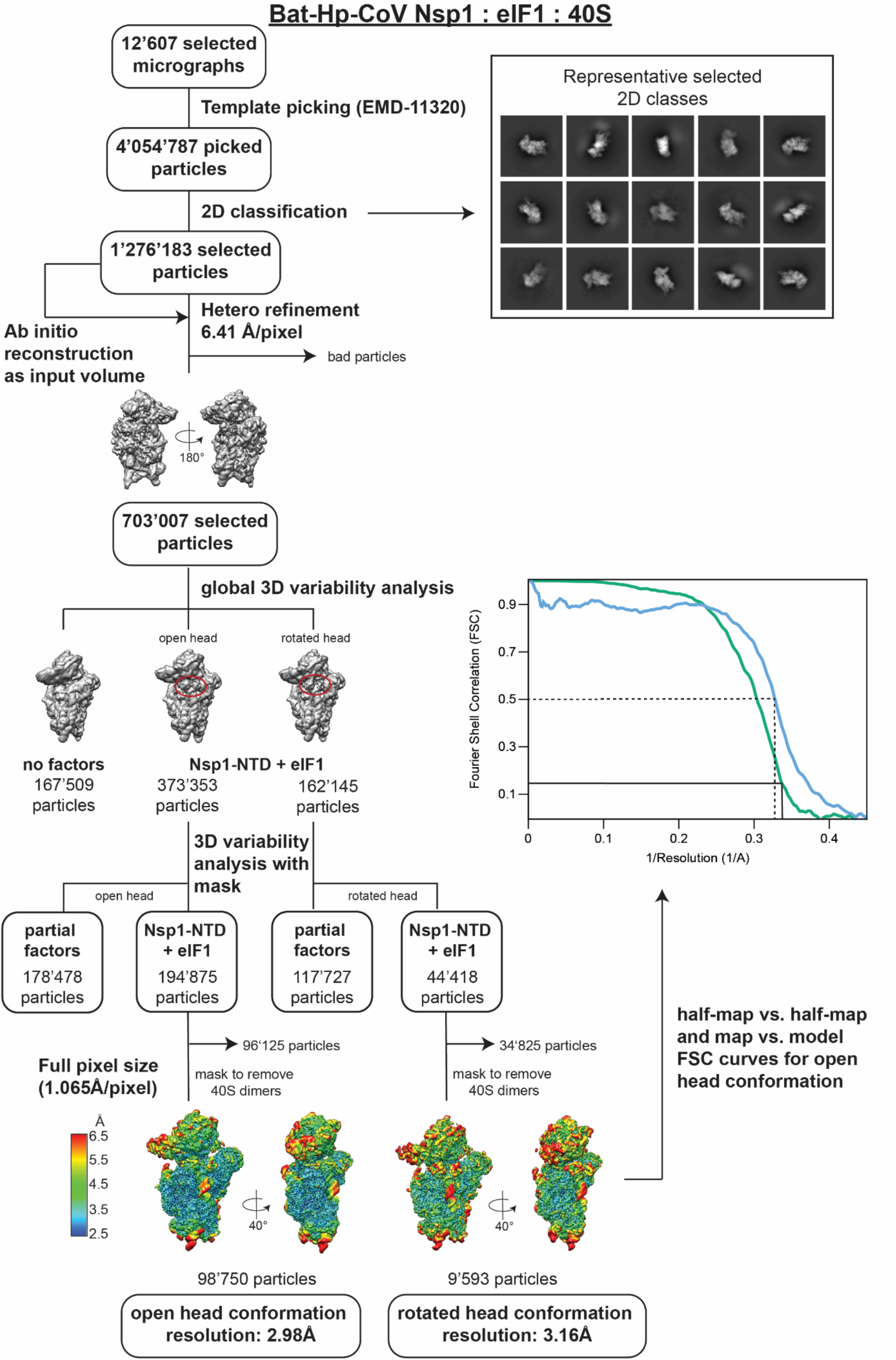
Scheme for the processing of the *in vitro* reconstituted Bat-Hp-CoV Nsp1 in complex with 40S and eIF1. Local resolution estimates are plotted as heat maps on the final volumes accompanied by the color scheme. The half map vs. half map and map. vs. model FSC curves are shown for the refinement.

**Figure S6:**
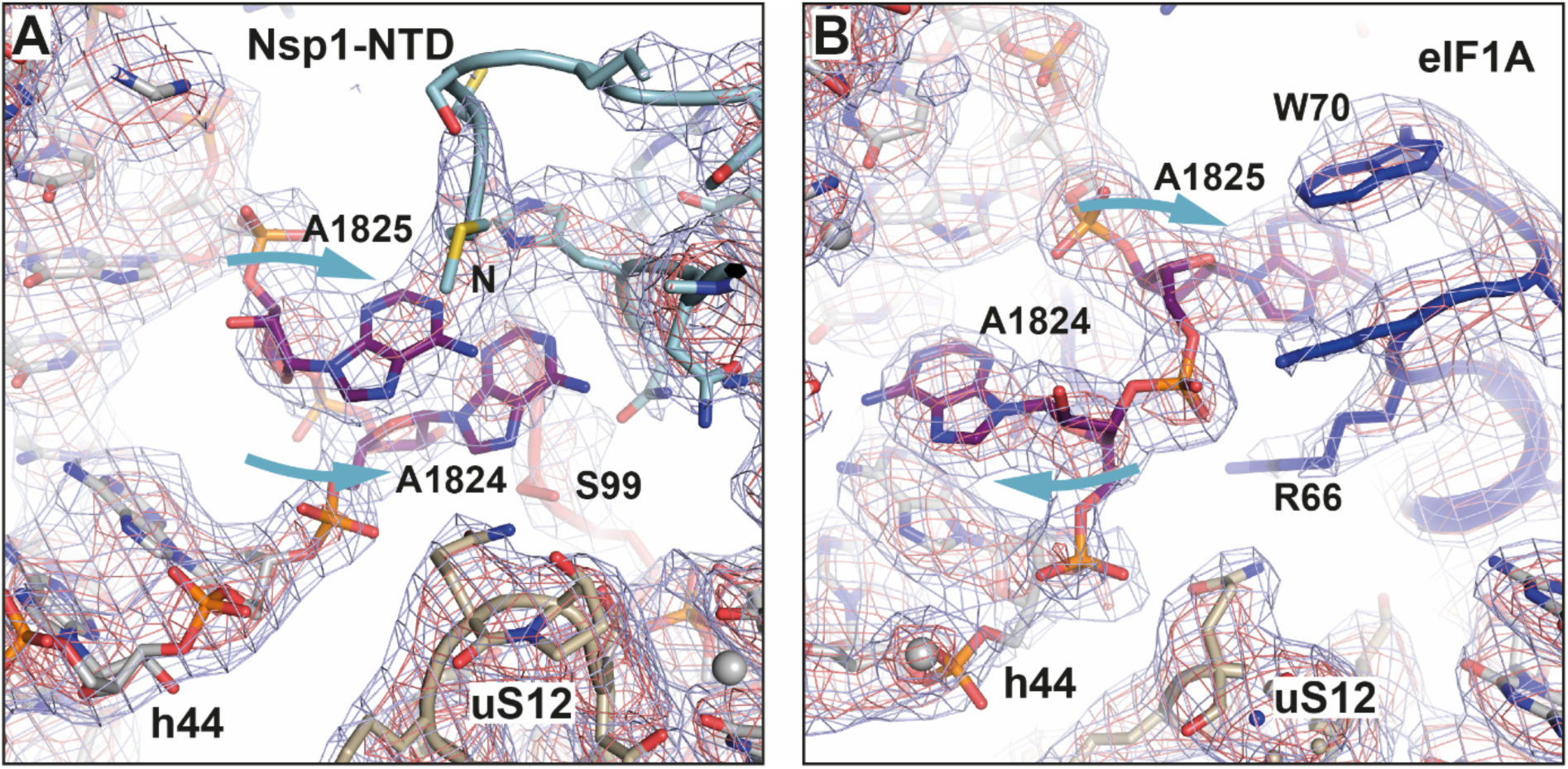
Comparison of decoding bases stabilized by either Bat-Hp-CoV Nsp1 or eIF1A. **(A)** Detailed view of the decoding site of the Bat-Hp-CoV Nsp1 – 40S – eIF1 complex. In the presence of the Nsp1 NTD, the decoding nucleotides A1824 and A1825 (purple) in h44 adopt a flipped-out conformation. The 3.0 Å experimental EM densities are shown as dark blue and red mesh and are contoured at 7*σ* and 11*σ*, respectively. **(B)** Detailed view of the decoding site of the MERS-CoV Nsp1 – 43S PIC. In the presence of eIF1A, decoding nucleotide A1825 adopts a flipped-out conformation, while A1824 remains flipped-in. The 2.65 Å experimental EM densities are shown as dark blue and red mesh and are contoured at 7*σ* and 11*σ*, respectively.

**Figure S7:**
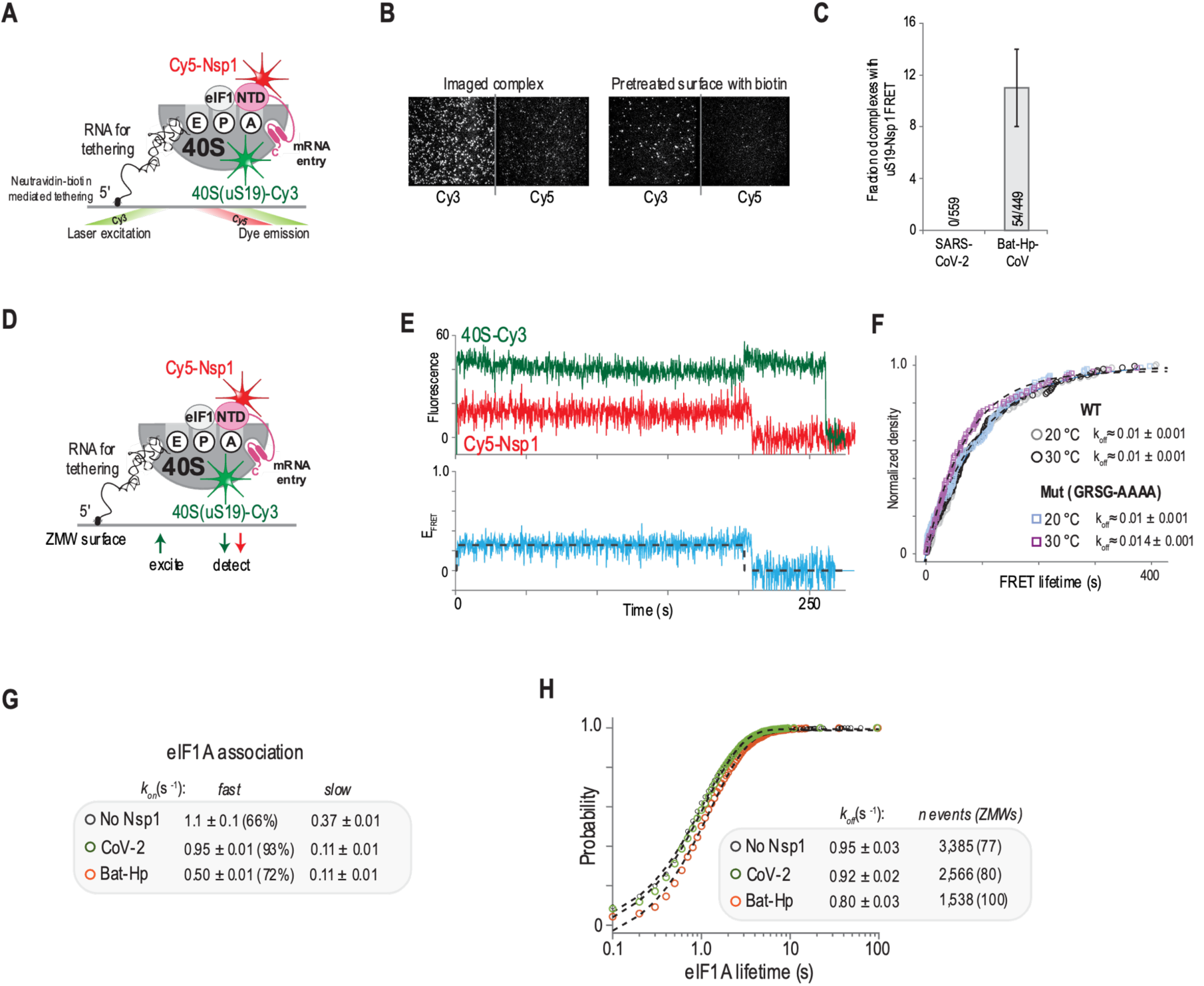
The NTD of Bat-Hp-CoV Nsp1 stably docks within the decoding center of the 40S subunit. **(A)** Schematic of the TIRFm single-molecule experiment setup performed at equilibrium at room temperature. The 40S subunit was labeled with Cy3 dye (FRET donor), and Bat-Hp-CoV Nsp1 with Cy5 dye (FRET acceptor). Fluorescence data were collected after direct excitation of the Cy3 dye via a 532 nm laser. **(B)** Example fields of view in the indicated conditions for the Bat-Hp-CoV Nsp1 complex, which indicates specific tethering of the complex. **(C)** Quantification of the fraction of tethered 40S complexes (scored via the Cy3 signal, green) that yielded FRET with either SARS-CoV-2 or Bat-Hp-CoV Nsp1 proteins (Cy5, red). The SARS-CoV-2 data is also reported in Figure 5C. **(D)** Schematic of the zero-mode-waveguide (ZMW) single-molecule experiment setup. The 40S subunit was labeled with Cy3 dye (FRET donor), and Nsp1 with Cy5 dye (FRET acceptor). Fluorescence data were collected after direct excitation of the Cy3 dye via a 532 nm laser. **(E)** Example single-molecule fluorescence data from a single ZMW for the Bat-Hp-CoV Nsp1(Cy5, red) – 40S (Cy3, green) complex imaged at equilibrium at 20 °C. **(F)** Plot of the observed FRET lifetimes observed for the indicated Bat-Hp-CoV Nsp1 proteins at the indicated temperatures. The dashed lines represent fits to an exponential function, which was used to derive the reported rates. **(G, H)** Quantification of eIF1A association times (panel G) and lifetimes (panel H) when the indicated Nsp1 proteins were present. Dashed lines represent fits to exponential functions. The data used to derive the eIF1A association rates are also reported in Figure 5F.

**Figure S8:**
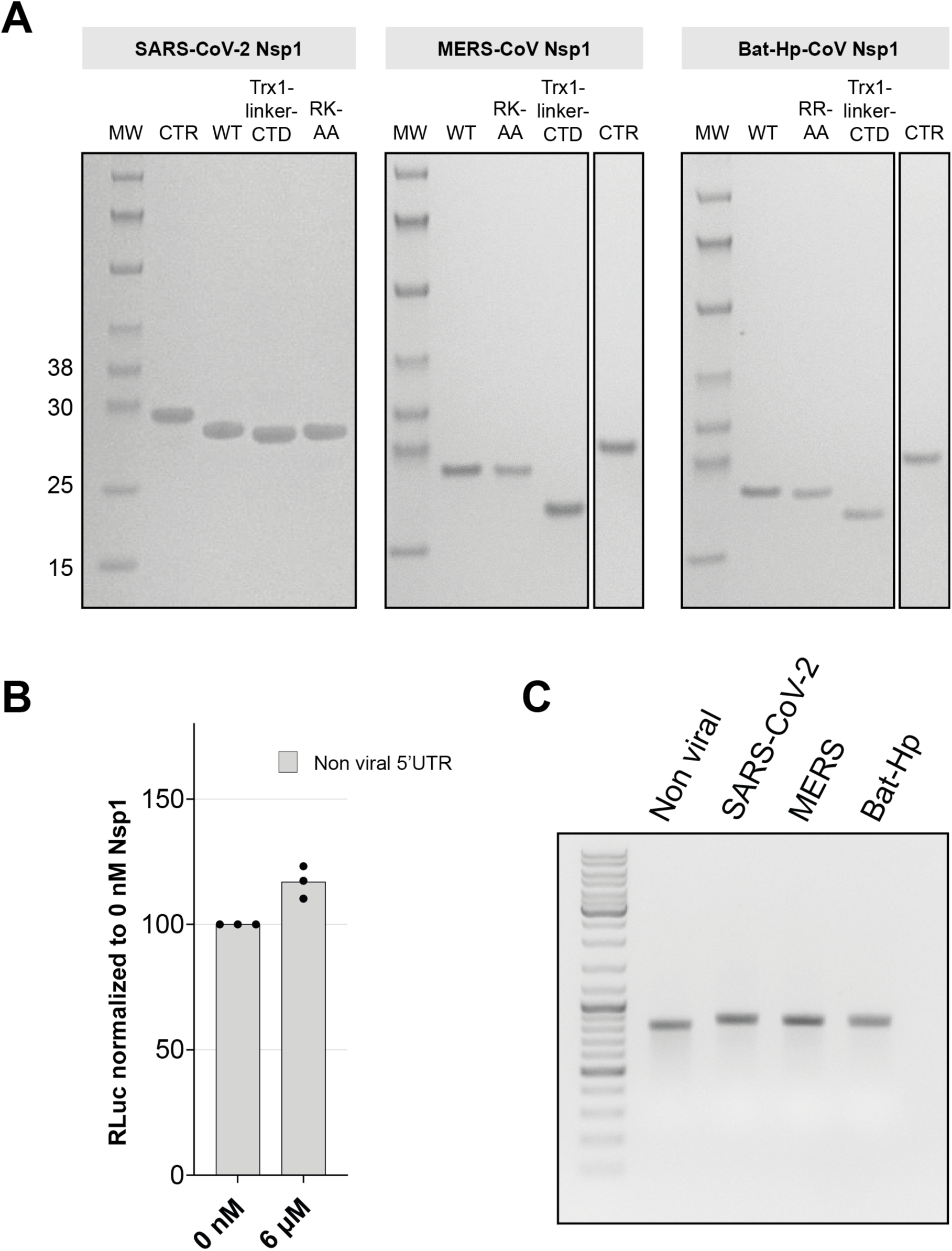
Components of the *in vitro* translation reaction. **(A)** Input purified protein samples for *in vitro* translation assays. Four μL of dialyzed Nsp1 protein samples at a concentration of 18 μM for SARS-CoV-2 and 9 μL of Bat-Hp-CoV and MERS-CoV Nsp1 were analyzed on a 4–12% SDS-PAGE and stained by Imperial protein staining (Thermo Fisher Scientific). **(B)** RLuc activity measurements of *in vitro* translation reactions containing 0 μM or 6 μM of purified recombinant control protein (Trx1-SSG-CTD KH SARS-CoV-2) normalized to the readout of the RLuc reporter mRNA in the absence of a control protein. Data are presented as mean values of three biological replicates (sets of translation reactions shown in points) averaged after three measurements. **(C)** 1% agarose gel electrophoresis of the *in vitro* transcribed RLuc reporters used for the *in vitro* translation assays.

## Notes

### Competing Interest Statement

The authors have declared no competing interest.

